# Emergent Electrical Activity, Tissue Heterogeneity, and Robustness in a Calcium Feedback Regulatory Model of the Sinoatrial Node

**DOI:** 10.1101/2022.11.11.516175

**Authors:** Nicolae Moise, Seth H. Weinberg

## Abstract

The sinoatrial node (SAN) is the primary pacemaker of the heart. SAN activity emerges at an early point in life and maintains a steady rhythm for the lifetime of the organism. The ion channel composition and currents of SAN cells can be influenced by a variety of factors. Therefore, the emergent activity and long-term stability imply some form of dynamical feedback control of SAN activity. We adapt a recent feedback model - previously utilized to describedion conductances in neurons - to a model of SAN cells and tissue. The model describes a minimal regulatory mechanism of ion channel conductances via feedback between intracellular calcium and an intrinsic target calcium level. By coupling a SAN cell to the calcium feedback model, we show that spontaneous electrical activity emerges from quiescence and is maintained at steady-state. In a 2D SAN tissue, spatial variability in intracellular calcium targets lead to significant, self-organized heterogeneous ion channel expression and calcium transients throughout the tissue. Further, multiple pacemaking regions appear, which interact and lead to time-varying cycle length, demonstrating that variability in heart rate is an emergent property of the feedback model. Finally, we demonstrate that the SAN tissue is robust to the silencing of leading cells or ion channel knockouts. Thus, the calcium feedback model can reproduce and explain many fundamental emergent properties of activity in the SAN that have been observed experimentally based on a minimal description of intracellular calcium and ion channel regulatory networks.

**Key Points Summary:**

- The robust function of the sinoatrial node (SAN) is reproduced in an intracellular calcium feedback model governing ion channel conductances
- The feedback model predicts the emergence and long-term maintenance of spontaneous oscillatory electrical activity
- Integrating the feedback model into a 2D SAN tissue leads to emergent spatial heterogeneity, multiple pacemaking regions, and variable cycle length
- SAN cells and tissue with feedback are robust to cell injury and channel knock-outs

## 1 Introduction

The sinoatrial node (SAN) is the primary pacemaker of the heart. The electrical impulses emerge from a population of spontaneous oscillating cells, which are inherently highly heterogeneous, with different sizes, ion channel conductances, and isolated spontaneous cycle lengths [1, 2, 3, 4, 5]. The SAN cells are organized into a network [6, 7] and are weakly coupled through gap junctions [8, 9]. Even with weak intercellular coupling, SAN cells synchronize and generate a coherent impulse of sufficient intensity to propagate to the atria and subsequently to the rest of the heart [10, 11, 12].

The SAN is an extremely robust system, which can maintain regular, repetitive spontaneous activity for very long periods of time (from the earliest developmental stages up to the entire lifetime of the organism), despite continuous ion channel turnover at the cellular level, heterogeneity between cells, and potential tissue remodeling or injury at the tissue level. In this study, we address several critical associated questions: How does this spontaneous activity emerge, and how is it maintained within physiological ranges for such long periods of time?

Recently, there has been a significant appreciation of the role of cardiac cell variability in theoretical models. This shift in thinking has led to consideration of populations of cells, each defined by different ion channel expression levels, and thus, broadly, variable phenotypes [13]. This variability is however constrained by the need to achieve ‘correct’ system outputs - in the case of the cardiac cell, a particular combination of ion channel expression levels are constrained to generate ‘good enough solutions’ (GES) for the time courses of the action potential and/or calcium transient [14, 15, 16]. The ability of the cells to both achieve and maintain a GES over time implies the existence of a feedback mechanism, adjusting subcellular characteristics (i.e. protein expression, channel post-translational modifications or overall membrane availability) to cell-level electrical activity. We posit that a feedback system can solve the two problems posed above: the emergence of ‘good enough’ activity, and its maintenance when subjected to perturbations.

Intracellular calcium concentration (*Ca*_*i*_) is a particularly fitting target for a feedback system, as has been investigated in a number of neuronal models [17, 18, 19]. While transmembrane voltage is the typical indicator of excitable cell electrical activity, it is difficult for a cell to sense and integrate voltage into regulatory networks. On the other hand, the calcium transient is by itself an important output of cellular electrical activity, especially in the SAN as part of the calcium clock driving automaticity [20, 21]. Further, *Ca*_*i*_ dynamics are tightly correlated with the transmembrane potential under normal conditions, such that *Ca*_*i*_ is an appropriate sensor of cellular electrical activity. Finally, *Ca*_*i*_ is an ubiquitous intracellular second messenger, implicated in a wide range of regulatory pathways, which can control protein expression or act directly on membrane channels to modulate their activity in neurons [22, 23], cardiomyocytes [24, 25, 26] and in particular in the SAN [27, 28].

Linking both GES and the role of calcium in the cardiac cell, Rees and colleagues [29] proposed a optimization-based protocol to find ‘correct’ ion channel conductance combinations in a ventricular cell model using *Ca*_*i*_ as a sensor. They showed that fitting only calcium transient targets (and not metrics based on the transmembrane potential) is sufficient for physiological action potential characteristics. Further, the optimization protocol imposed fixed ratios between conductances, in particular between inward calcium currents and outward potassium currents. The ‘good enough solutions’ that emerge from the optimization protocol can therefore be thought as the preferred, physiological steady-states of channel expression in the cardiac cells. While providing significant insight into cardiac cellular variability, this approach is limited by the fact that the model does not consider the dynamics of ion conductances - specifically, how does the cell reach and maintain this preferred state, and how does the cell respond to perturbations?

Such questions have similarly been explored in the computational neuroscience field. In one such seminal study, O’Leary and colleagues assumed that such GES (expressed as a particular ratio between ion channel conductances) exist for particular types of neurons [30]. The authors proposed a minimal dynamical feedback model through which neurons can generate and maintain their particular ion channel conductance composition. Briefly, the cell tries to ‘match’ some set intracellular calcium target by regulating its ion channel mRNA and thus ion channel membrane expression levels. If *Ca*_*i*_ is below the target, ion channel expression increases, thus leading to higher average *Ca*_*i*_, while conversely, when *Ca*_*i*_ is above the target, this leads to a decrease in ion channel conductance. Crucially, the ratio between channel conductances is always strictly maintained. Using this feedback model, the authors showed that electrical activity emerges and is then sustained once the intracellular calcium reaches the target level. By varying different channel ratios, different neuronal phenotypes (such as continuously oscillating or various bursting patterns) emerge. Further, the feedback model determined the emergence of activity in simple oscillatory networks in a manner robust to injury. Finally, cells responded to channel knockouts (KOs) in both an adaptive and pathological manner, depending on the targeted channel.

In this study, we adapt the minimal feedback model proposed by O’Leary et al to the rabbit SAN cell (as described by Severi and colleagues [31]) and further extend this approach to model SAN tissue. We find that, in the single cell, sustained spontaneous electrical activity emerges from a low-conductances quiescent initial state. Spontaneous activity is maintained over long periods of time, and is adaptable and robust to changes in the internal *Ca*_*i*_ target, external pacing, and heterogeneity in feedback model parameters. In SAN tissue, coupled cells with heterogeneous calcium targets self-organize into a spatially heterogeneous ion channel conductance expression pattern, leading to the emergence of multiple pacemaking sites and an inherently variable tissue-level cycle length. Finally, we show that the SAN tissue model with feedback is robust to injury or loss (i.e., KOs) of key pacemaking ion channels.

## 2 Methods

### Single SAN cell model

The basis of our study is a model of the rabbit SAN cell described by Severi and colleagues [31]. The transmembrane potential *V* dynamics are governed by

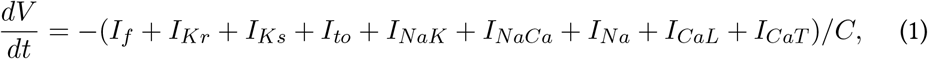

where *C* is the membrane capacitance, and *I*_*x*_ (where *x ∈* {*f, Kr, Ks, to, NaK, NaCa, Na, CaL, CaT*}) denote the currents carried by the sarcolemmal ion channels, pumps, and exchangers. For compactness, we can write Eqn. 1 as

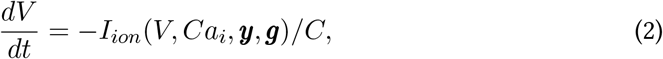

where we have denoted that the sum of the ionic currents can be written as a function of *V*, intracellular calcium (*Ca*_*i*_), ***y*** (a vector collecting gating variables, state variables, and intracellular compartment ionic concentrations), and ***g*** (a vector collecting the conductances for the ionic currents). In typical excitable cell models, including the baseline Severi et al. SAN cell model, conductances ***g*** are defined as fixed constant values; however, via the feedback model (as described below), these conductances are time-varying state variables.

The dynamics of ***y*** are governed by

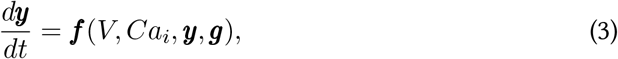

where the function ***f*** accounts for the gating variable, state variable, and concentration dynamics. Most of the membrane currents follow standard Hodgkin-Huxley formulations (*I*_*f*_, *I*_*CaT*_, *I*_*CaL*_, *I*_*Kr*_, *I*_*Ks*_, *I*_*to*_), with kinetics fitted to recent experimental data. Critically for our study, the model also incorporates a detailed description of *Ca*_*i*_ dynamics (derived from the Maltsev-Lakatta model [32]), which includes SERCA uptake of cytoplasmic Ca^2+^ into the sarcoplasmic reticulum (SR), flow of Ca^2+^ between network SR and junctional SR, and release by RyR into a sub-sarcolemmal space. Again, for compactness, we can write *Ca*_*i*_ dynamics as

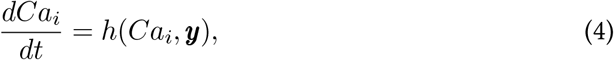

where function *h* describes how *Ca*_*i*_ dynamics depend on *Ca*_*i*_ and other calcium handling fluxes. The full details of gating variable, state variable, and ion concentration dynamics, and associated parameters (as described by ***f*** and *h*) are provided in Severi et al [31].

### Calcium feedback model

To describe regulation of the conductances, we adapted the O’Leary feedback model [30] of neuronal channel conductance regulation to the SAN cell. The fundamental assumption of the O’Leary model is that cells have an inherent calcium target (*Ca*_*tgt*_) for intracellular calcium *Ca*_*i*_ levels (with *Ca*_*i*_ dynamics described by the Severi model in this study), such that *Ca*_*i*_ levels are driven to “match” the target via regulation of ionic channel expression and thus conductances. The feedback model is expressed by

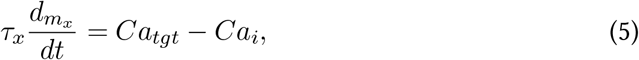

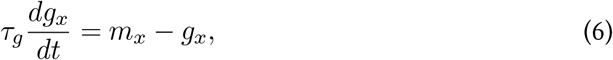

where conductance vector ***g*** contains all *g*_*x*_ (for *x* noted above, with an additional term for the SERCA flux).

The feedback model and associated variables can be explained as follows: The difference between the cellular *Ca*_*tgt*_ and actual *Ca*_*i*_ level drives the dynamics of *m*_*x*_, where *m*_*x*_ is an intermediary between the sensing of calcium differences and the expression of conductances and can broadly represent a term proportional to the mRNA level corresponding to the particular channel *x*. The dynamics of *m*_*x*_ drives subsequent changes in the conductance *g*_*x*_. The dynamics are governed by time constants *τ*_*x*_, which is specific for each species *x*, and *τ*_*g*_, which is equal for all cases. Importantly, the ratio between each *τ*_*x*_ sets the conductance ratio between the channels, which is always maintained (see Figure 1). The values for *τ*_*x*_ are given in Table S1. Note that *τP*_*up*_ is negative, as we assumed that SERCA decreased its activity if *Ca*_*i*_ *< Ca*_*tgt*_.

**Figure 1.**
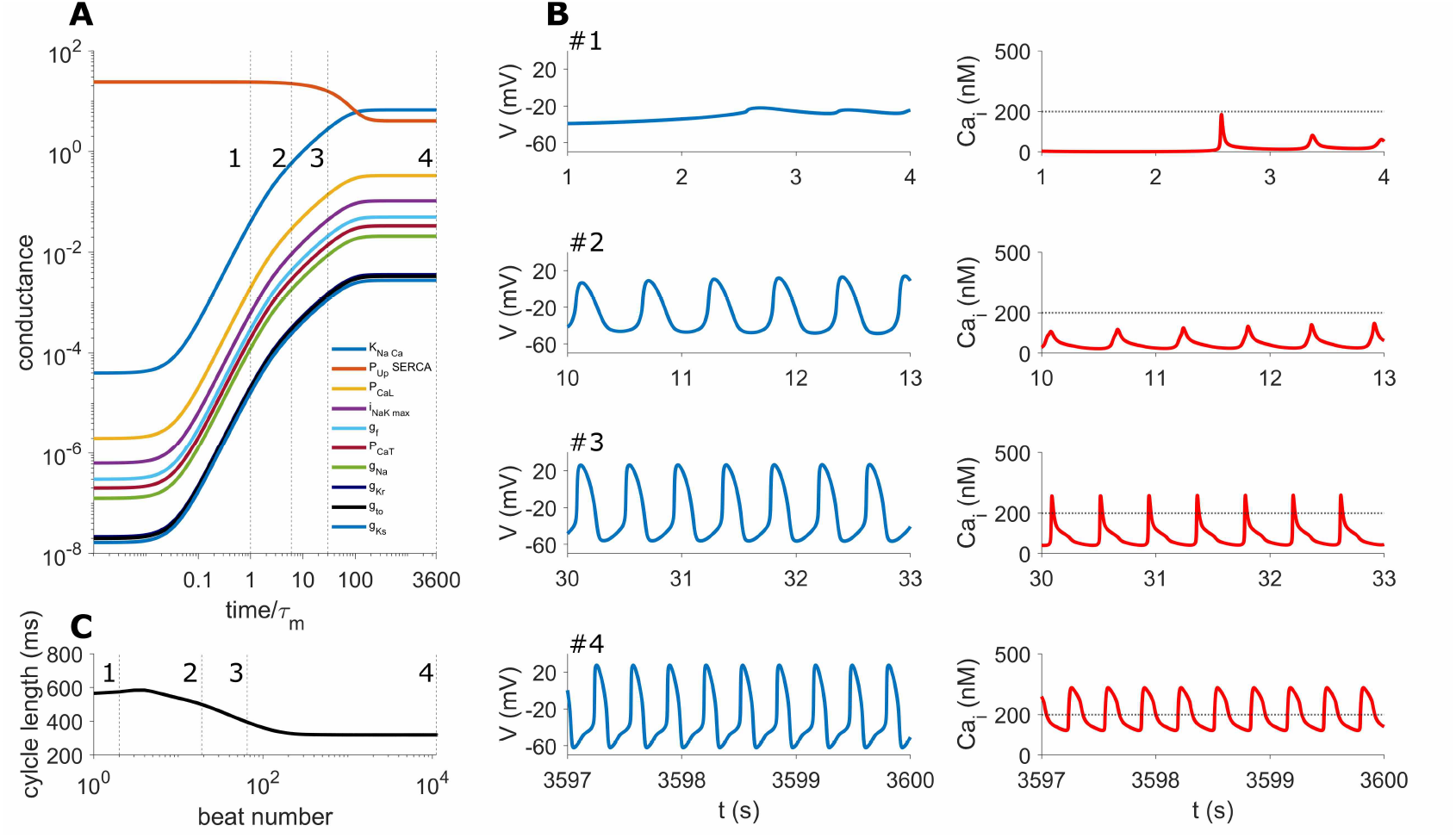
Emergent spontaneous activity in single SAN cells **A**. Conductance values generally increase (except SERCA) and approach a steady-state level, as spontaneous rhythmic activity emerges. All conductances have units of *μ*S, except *K*_*NaCa*_, *I*_*NaK max*_ (nA), *P*_*CaT*_, *P*_*CaL*_ (nA/mM) and *P*_*up*_ (mM/s). **B**. Voltage and *Ca*_*i*_ traces at different times of the simulation. The horizontal dotted line in the calcium plots marks the set *Ca*_*tgt*_ = 200 nM. **C**. Cycle length of spontaneous activity evolution over time. The vertical lines in **A** and **C** denotes the times highlighted in **B**.

### 2D SAN tissue model

Extending the single cell model to a 2D tissue, the voltage dynamics are described by the following partial differential equation

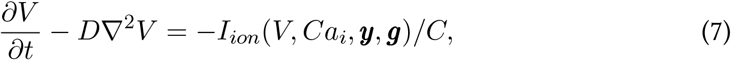

where *D* is the diffusion coefficient governing cell-cell coupling, and variables *V, Ca*_*i*_, ***y***, and ***g*** are functions of space (in addition to time).

### Calcium feedback model time scale

We empirically found that the SAN tissue required a longer time to reach steady state, compared with the single cell. Therefore, due to computational time constraints, we increased the sensitivity of the feedback model by decreasing all *τ*_*x*_ parameters by a factor of 10 (see Table S1). We denote the relative time scaling factor by *τ*_*m*_, taking therefore *τ*_*m*_ = 1 in the single cell case and *τ*_*m*_ = 0.1 in the tissue simulations. Accordingly, we plot all the conductance time courses relative to this time scale *τ*_*m*_.

Finally, an important assumption of the model is that the conductance changes occur on a significantly slower time scale than the dynamics of the membrane potential and *Ca*_*i*_ changes. We show that this assumption is valid for the time constant parameter choices of both the single cell and tissue simulations (Figure S1). Although conductances oscillate in coordination with the calcium transient (CT), the amplitude of the conductance oscillations are insignificantly small on the time scale of the action potential (AP) or CT.

### Numerical methods

Single cell simulations were integrated using the forward Euler method, with a time step of 0.1 ms. To decrease computational time, we use an operator splitting method for the intracellular calcium dynamics (the equations for SR Ca^2+^ release, *I*_*NaCa*_ and Ca^2+^ fluxes between intracellular compartments, as described in Severi et al [31]), due to increased numerical stiffness compared to the rest of the system. To prevent the model from representing non-physiological conditions, we imposed a threshold to all ionic conductances *g*_*x*_, such that all membrane conductances can vary between 1*/*40 and 40 times their baseline values (as given in Severi et al [31]), while *P*_*up SERCA*_ can vary between 1*/*10 and 10 times its baseline value.

In the 2D SAN tissue, the spatial domain is a 20 × 20 grid, with a spatial discretization Δ = 0.0125 cm, which is roughly the size of the SAN cell, such that each spatial grid location corresponds with a single SAN cell. We use an operator splitting to integrate the voltage diffusion term and the rest of the system. The voltage diffusion term is solved using the alternating-direction implicit (ADI) method, while the remaining variables are integrated with forward Euler, both using a time step of 0.1 ms.

### SAN cell and tissue perturbations

Channel knockout (KO) are represented by scaling the conductance of the channel with a value less than 1 (based on the degree of KO) and fixing the conductance at this smaller value, such that the conductance level is decoupled from the feedback model. All the other conductances remain variable and adapt to the changes induced via the feedback model. For the cell ablation experiments in SAN tissue, the *Ca*_*tgt*_ value for the cells with the 20 highest conductance values are set to 0 (described further below).

## 3 Results

### Emergent spontaneous activity in a single SAN cell

We start by investigating the emergence of electrical activity in the single SAN cell. The initial values for ion channel conductances are close to zero for all species except SERCA (Fig 1A), consistent with a quiescent cell, i.e., there is no rhythmic electrical activity and no calcium transients (CTs) (Fig 1B, #1). As a result, *Ca*_*i*_ is below the set *Ca*_*tgt*_, promoting an increase in ionic current conductances, which gradually leads to incipient rhythmic activity (Fig 1B, #2), followed by the acceleration of the rhythm (i.e., a decrease in the spontaneous cycle length, Fig 1C). In parallel, the amplitudes of the APs and the CTs also increase. When *Ca*_*i*_ reaches a sufficient amplitude and frequency (Fig 1, #4), the conductance values plateau, and the cell reaches its stable equilibrium activity and a constant cycle length. Thus, we find that this minimal feedback model is sufficient to develop and maintain normal spontaneous electrical activity, from low minimal ion channel expression levels. We note that conductance levels always maintain the same ratio between each other, as set by the time constants in the feedback model (see Methods). Therefore, for simplicity in future figures, we only present the dynamics of *g*_*Na*_, as the dynamics of one conductance level is representative of all of the conductances.

We next consider the role of the *Ca*_*tgt*_ value on the SAN cell equilibrium. We find that varying *Ca*_*tgt*_ results in different steady-state conductance levels and spontaneous cycle lengths (Fig 2A, B). Increasing *Ca*_*tgt*_ leads to an increase in steady-state conductance level, which in turn is associated with faster spontaneous electrical activity (i.e., decreased cycle length). Moreover, the higher conductances promote changes in the morphology of the APs and CTs, with generally higher amplitudes (Fig 2C, #1-3).

**Figure 2.**
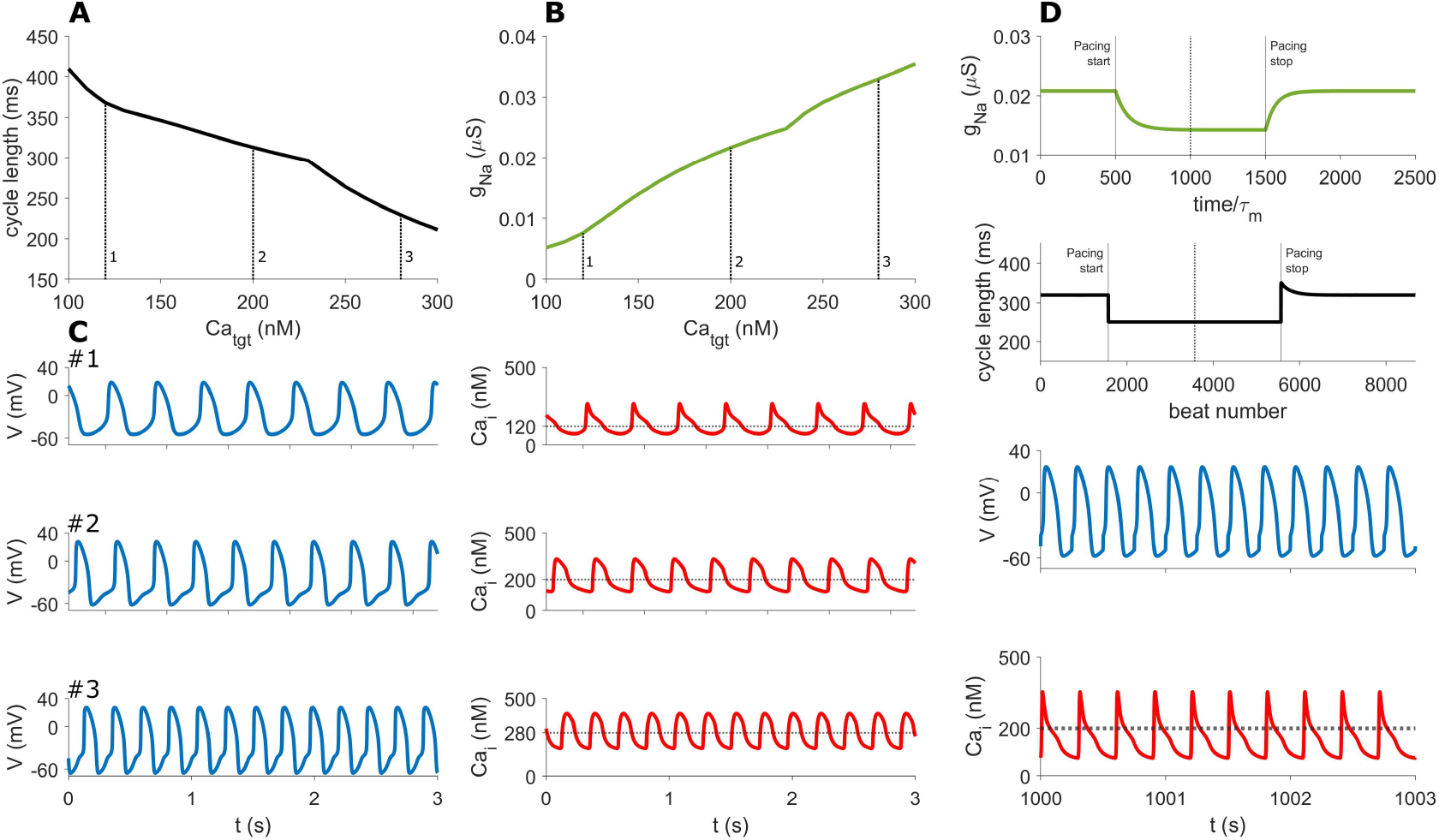
Calcium target sets spontaneous activity cycle length and ion channel conductance levels in SAN cells. Increasing *Ca*_*tgt*_ (**A**) decreases spontaneous activity cycle length and (**B**) increased conductances. **C**. Voltage and *Ca*_*i*_ activity after reaching steady-state for 3 different *Ca*_*tgt*_ values, denoted in **A** and **B. D**. (Top) Conductance and cycle length are shown as a function of beat number before and after external pacing at a faster rate than the intrinsic rate. This pacing leads to a compensatory decrease in conductances and thus a smaller calcium transient. The vertical dotted line denote the time for the (Bottom) voltage and intracellular calcium traces.

The changes in both cycle length and CT morphology highlight that the *Ca*_*tgt*_ can thus be “met” through two (correlated) approaches - regulation of the CT amplitude and changing spontaneous activity frequency. We therefore next consider the cellular response in the presence of direct control of the frequency - specifically in response to externally controlling frequency through external pacing. Starting from the equilibrium state for *Ca*_*tgt*_ = 200 nM, we then externally pace the SAN cell at a cycle length faster than the equilibrium cycle length (Fig 2D). Due to the faster activity, the conductances decrease, until a new equilibrium is reached. The frequency however cannot change as it is being imposed externally. However, lower conductances lead instead to a smaller CT, as an adaptation to the faster pacing rate (c.f., Fig. 2C, #2 and D). Following termination of external pacing, the conductances return to the previous high conductance levels. However, between the end of pacing and the return to steady-state, there is a transient period of slow spontaneous activity (i.e., longer cycle length), due to the remodeled lower conductance levels.

As an additional demonstration of the feedback model robustness, we perform a population study comprised of 1000 simulations for a fixed *Ca*_*tgt*_ of 200 nM. In each simulation, the feedback model time constants *τ*_*x*_ for each ionic current are scaled by random value uniformly distributed between 0.5 to 2, and the steady-state conductance levels and cycle lengths are measured. Importantly, 998 out of the 1000 time constant combinations resulted in spontaneous electrical activity in the SAN cell. Histograms of the steady-state *g*_*Na*_ and cycle length, shown in Fig. S2, demonstrate the variability in ionic conductances and spontaneous activity. Critically, these simulations demonstrate 1) that SAN spontaneous electrical activity is robust to changes in the time constants, and thus changes in the time constant ratios, in the feedback model, and 2) that variability in spontaneous activity and conductance levels can be achieved through variation in either the target calcium target or the feedback model time constants. Going forward, for simplicity we introduce heterogeneity in SAN cell activity by varying *Ca*_*tgt*_; however, in a physiological setting, variability in individual SAN cell properties is likely a combination of heterogeneity in ion channel expression signaling (as represented by the feedback model time constants *τ*_*x*_) and calcium signaling (as represented by *Ca*_*tgt*_).

### Emergent spontaneous activity and variability in a 2D SAN tissue

We next extend the feedback model to a 2D SAN tissue (Fig. 3), consisting of a 20 × 20 array of cells. We represent spatial heterogeneity in the SAN tissue, such that each cell has a different *Ca*_*tgt*_, drawn from a uniform distribution between 100 and 300 nM (Fig. 3A). As described in the Methods, cells are electrically coupled, with diffusion term *D* (constant throughout the tissue), representing gap junctional coupling. As in the single cell simulations, all conductances are initially low, consistent with quiescent cells. Conductance levels, as represented by *g*_*Na*_, for all 400 cells are shown as a function of time (Fig. 3C, Movie S1). We find that in some cells, *g*_*Na*_ increase in time, while in others, *g*_*Na*_ remains quite low, even decreasing compared with initial levels. In all cells, *g*_*Na*_ reaches an equilibrium level, with the steady-state spatial pattern of conductance values shown in Fig. 3B. The final *g*_*Na*_ distribution that emerges is very wide (Fig. 3D): most of the cells have low conductance values, with a minority of cells having very high values. Note also that the *g*_*Na*_ range is much wider in the SAN tissue, compared with the range from single cells with the same *Ca*_*tgt*_ values (cf. Fig. 2A, also see Fig. S3).

**Figure 3.**
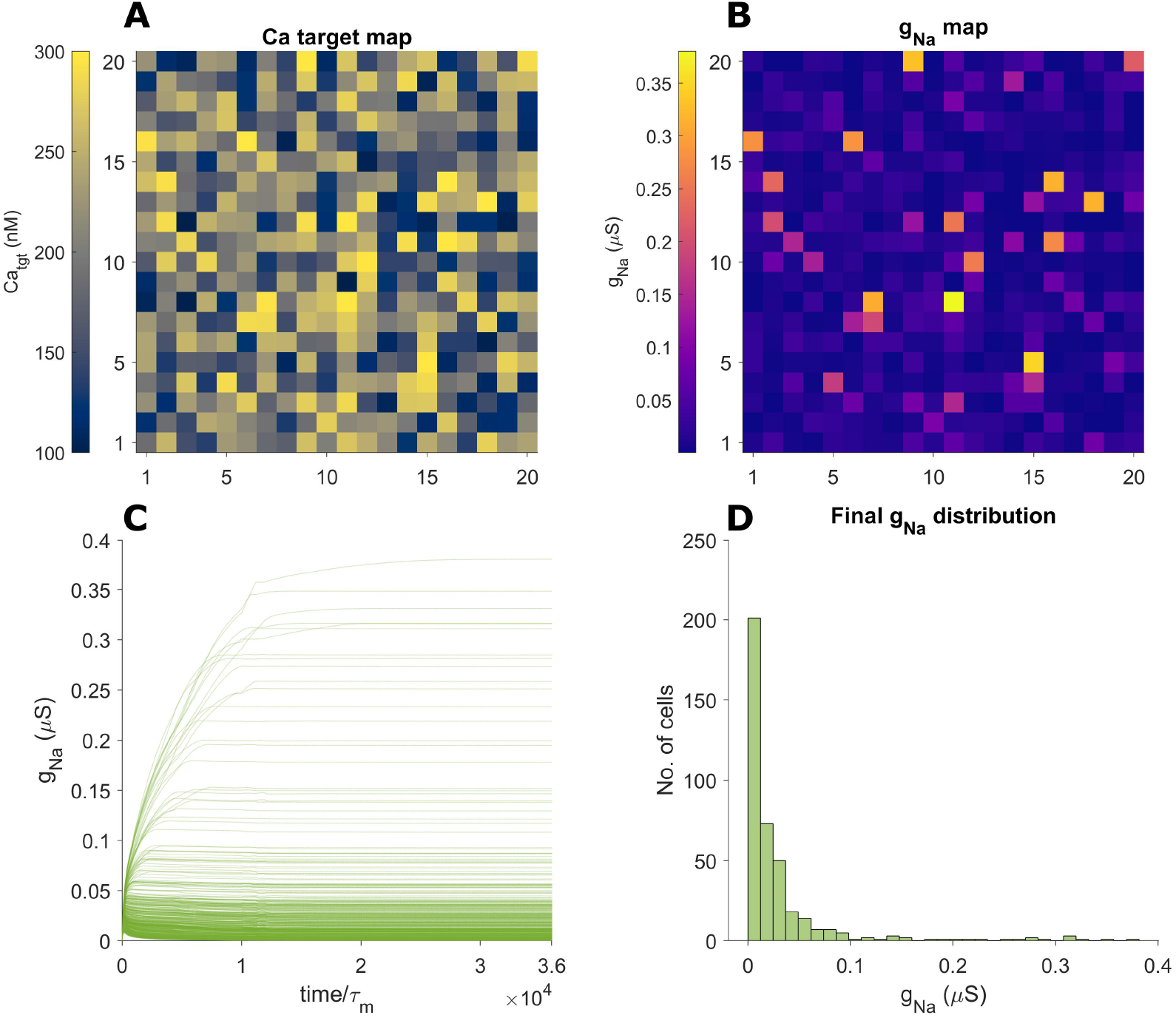
Distribution of conductances in a 2D SAN tissue. **A**. Spatial map of the random *Ca*_*tgt*_ values assigned to each cell in the SAN tissue. **B**. Spatial map of sodium conductance levels (*g*_*Na*_) after reaching steady-state. **C**. Evolution of conductances over time for all 400 cells in the SAN tissue. **D**. Final distribution of conductances in the tissue.

We next investigate the dynamics of the spontaneous activity that emerges from the SAN tissue. The highly heterogeneous conductance distribution leads to a variable tissue cycle length. When spontaneous activity first emerges, the cycle length is initially relatively long, but accelerates as conductances increase (Fig. 4A) and settles into a variable pattern, specifically of five different repeating cycle lengths, as shown in Fig. 4B. Interestingly, the average peak *Ca*_*i*_ also differs between beats (Fig. 4D). The AP morphology and duration is very similar between cells in the tissue, due to the cell-cell coupling (Fig. 4E). However, the CTs are heterogeneous, with a wide range of peak values and variable activation time (Fig. 4F, see also Fig. 5). Investigation of the voltage activation maps for these 5 beats (Fig. 4G, Movie S2) reveals that spontaneous activity emerges from multiple spatial locations, which alternate as the leading pacemaker of the 2D SAN tissue. Comparing the *g*_*Na*_ spatial pattern (cf. Fig. 3B) further reveals that the foci of these leading pacemaker sites correspond to the spatial location of regions of high conductance “clusters”. Note that the activation sequences in beat #3 and #5 are different, even though they initiate with the same cluster, likely due to the different activation pattern on the previous beat.

**Figure 4.**
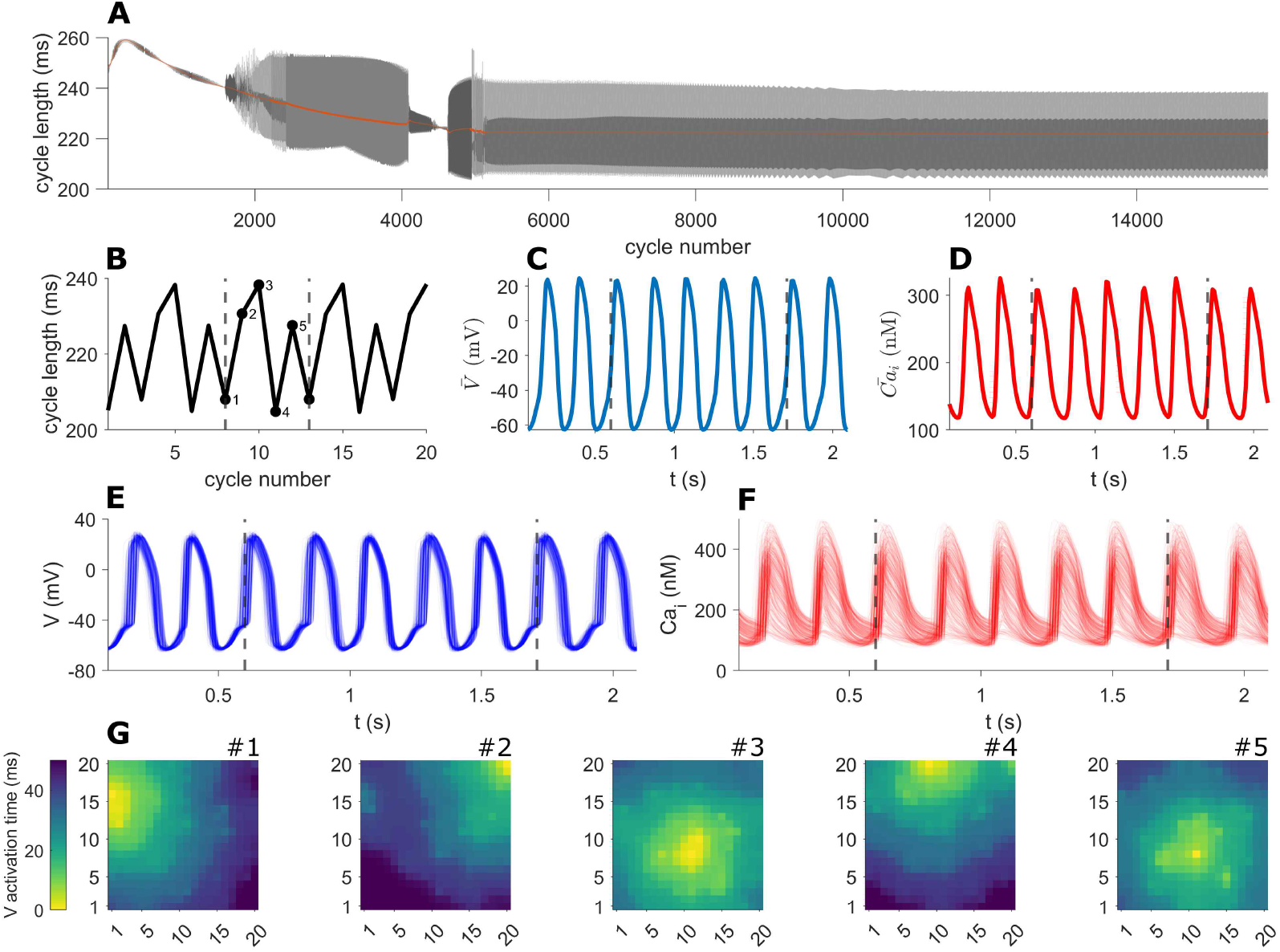
Emergent spontaneous activity with variable cycle length and multiple pacemaking sites in a 2D SAN tissue. **A**. Cycle length of the SAN tissue (gray), calculated from the peak-to-peak difference of the average voltage across the tissue. Red line denotes a running average of the cycle length to illustrate average trends. **B**. Highlight of steadystate cycle length pattern, consisting of 5 repeating cycle length values. **C**. Average tissue voltage and **D**. average *Ca*_*i*_ at steady state. The dotted vertical lines mark the repeating sequence of activating patterns from **B. E**. Voltage and **F**. *Ca*_*i*_ traces for all 400 cells in the 2D SAN tissue. **G**. Voltage activation maps for the 5 different activation patterns illustrates the multiple pacemaking sites.

**Figure 5.**
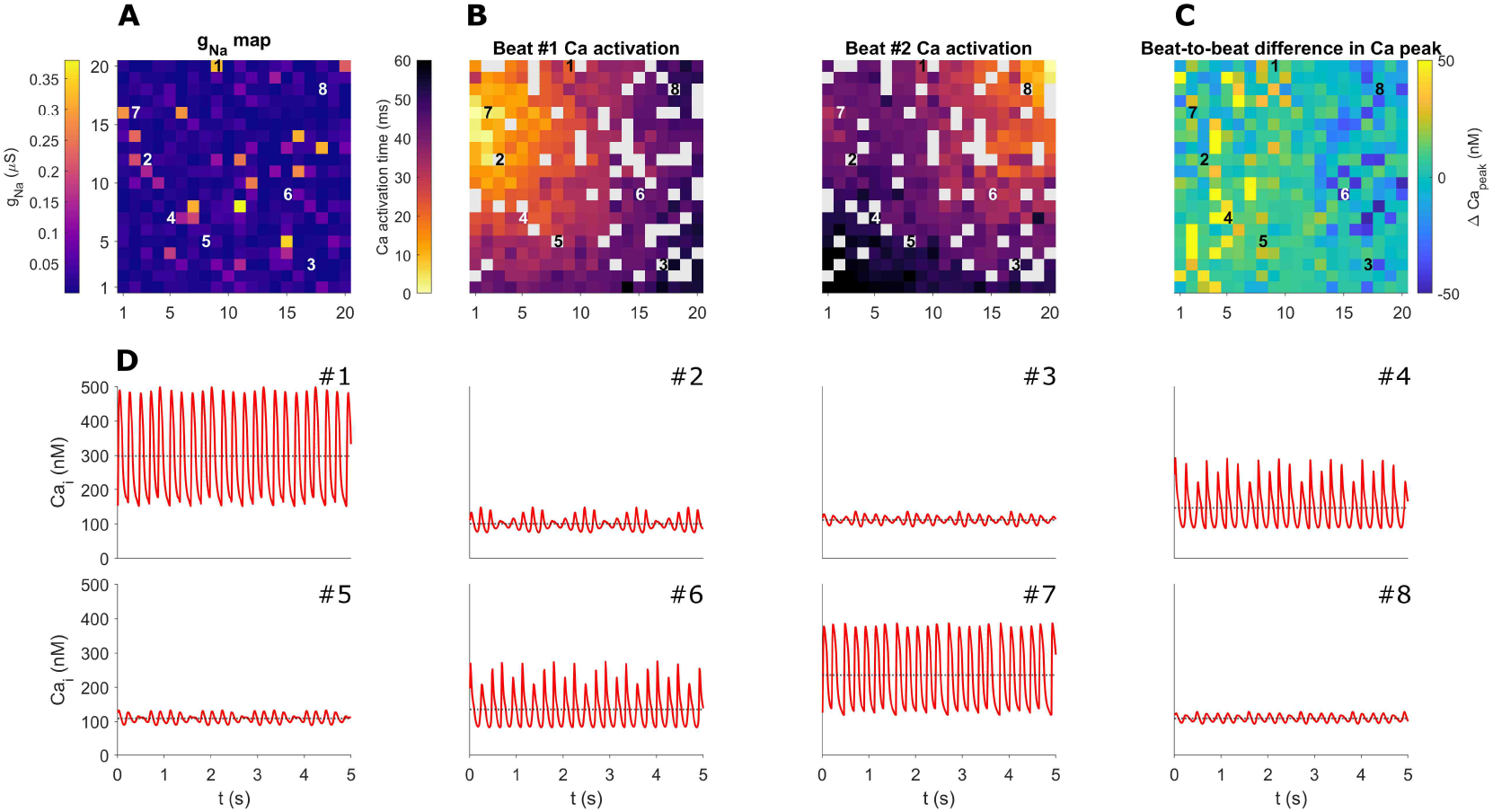
Heterogeneous *Ca*_*i*_ dynamics in the SAN tissue. **A**. Spatial map of sodium conductances (*g*_*Na*_) at steady state. **B**. *Ca*_*i*_ activation maps, corresponding to beats #1 and #2 in Fig. 4. The cell is considered to be activated above a threshold of 200 nM. The grey pixels represent cells with peak calcium levels below this value on a given beat. **C**. Beat-to-beat differences in calcium transient peaks, between the two beats shown in **B. D**. Example *Ca*_*i*_ traces from the SAN tissue, illustrating heterogeneity in calcium transient characteristics. The horizontal dotted lines show the associated *Ca*_*tgt*_ for that cell. The corresponding cells are numbered in **A**-**C**.

We next consider the spatial relationship between conductance levels and CT heterogeneity (Fig. 5). The *g*_*Na*_ spatial map (from Fig. 3B) is shown here for ease of comparison in Fig. 5A, with 8 spatial locations identified. The CT activation map from beats #1 and #2 (corresponding to the first two beats in Fig. 4G) illustrate that activation of calcium is less synchronized compared with voltage, with some cells activating slower than their neighbors (Fig. 5B). Some cells, shown as grey pixels, have low-amplitude CTs, below the (arbitrary) threshold of 200 nM. Notably, some cells cross the threshold on one beat but not the other. This can also be observed in Fig. 5C, which illustrates that there is considerable beat-to-beat differences between CT peaks. In Fig. 5D, we present example *Ca*_*i*_ traces from the 8 identified locations in the 2D tissue. Again, CT heterogeneity is readily apparent in these examples - there is a wide range of phenotypes, with some cells exhibiting small oscillatory activity (#2, 3, 5, 8), while others have high amplitude CTs (#1, 7). There are also complex peak CT dynamics, with some cells (in particular, #2, 4, 6) illustrating alternating patterns. Note that the dynamics follow the five-beat cycle highlighted in Fig. 4. The heterogeneity is primarily driven by each cells’ respective *Ca*_*tgt*_ (shown as the horizontal dotted line).

The SAN tissue results illustrate that the dynamic feedback control of ion channel conductances leads to highly heterogeneous conductance expression in tissue, as determined by the spatial variability in the cell’s respective *Ca*_*tgt*_ levels. Following the emergence of spontaneous activity, ion channel conductance levels reach steady-state values, with a distribution much wider than observed in isolated cells (for the same range of *Ca*_*tgt*_ values). At equilibrium, spontaneous electrical activation emerges from a number of cell clusters, which alternate as the leading pacemakers with no cluster becoming dominant over long periods of time. The spatial variation of impulse emergence leads to an inherently variable tissue-level cycle length. Underlying the complex voltage dynamics is highly heterogeneous calcium activity, which is a direct result of the feedback model. The heterogeneity in CTs and multifocal emergence of spontaneous activity agree quite closely with recent experimental data on calcium activity in intact SAN tissue [7] (see Discussion).

The prior results demonstrate the emergence of spontaneous activity with emergent cycle length variability and multiple leading pacemaking sites, due to *Ca*_*tgt*_ spatial heterogeneity, in the SAN tissue feedback model, for specific values for the *Ca*_*tgt*_ variability and electrical coupling diffusion coefficient *D*. We next demonstrate the generality of these results by systematically varying the two tissue-level parameters, diffusion coefficient *D* and the range of the random distribution of *Ca*_*tgt*_ values, Δ*Ca*_*tgt*_ (Fig. 6). For a range of values of both parameters, we measure channel conductances heterogeneity (Δ*g*_*Na*_, defined as the difference of the maximum and minimum steady-state *g*_*Na*_ values across the tissue) and cycle length heterogeneity (*σ*, defined as the cycle length standard deviation once the equilibrium is reached). We find that above a minimum degree of *Ca*_*tgt*_ variability, specifically Δ*Ca*_*tgt*_ = 60 nM, the conductance values are highly variable (Fig. 6A). The heterogeneity in channel conductances is accompanied by a corresponding variability in cycle length (Fig. 6B). However, above this threshold, increasing diffusion coefficient *D* tends to decrease cycle length variability. That is, increased tissue coupling promotes synchronization between the different emerging pacemaking clusters and therefore leads to less variable tissue cycle length. Increased *D* also leads to a small decrease in Δ*g*_*Na*_ (Fig S3).

**Figure 6.**
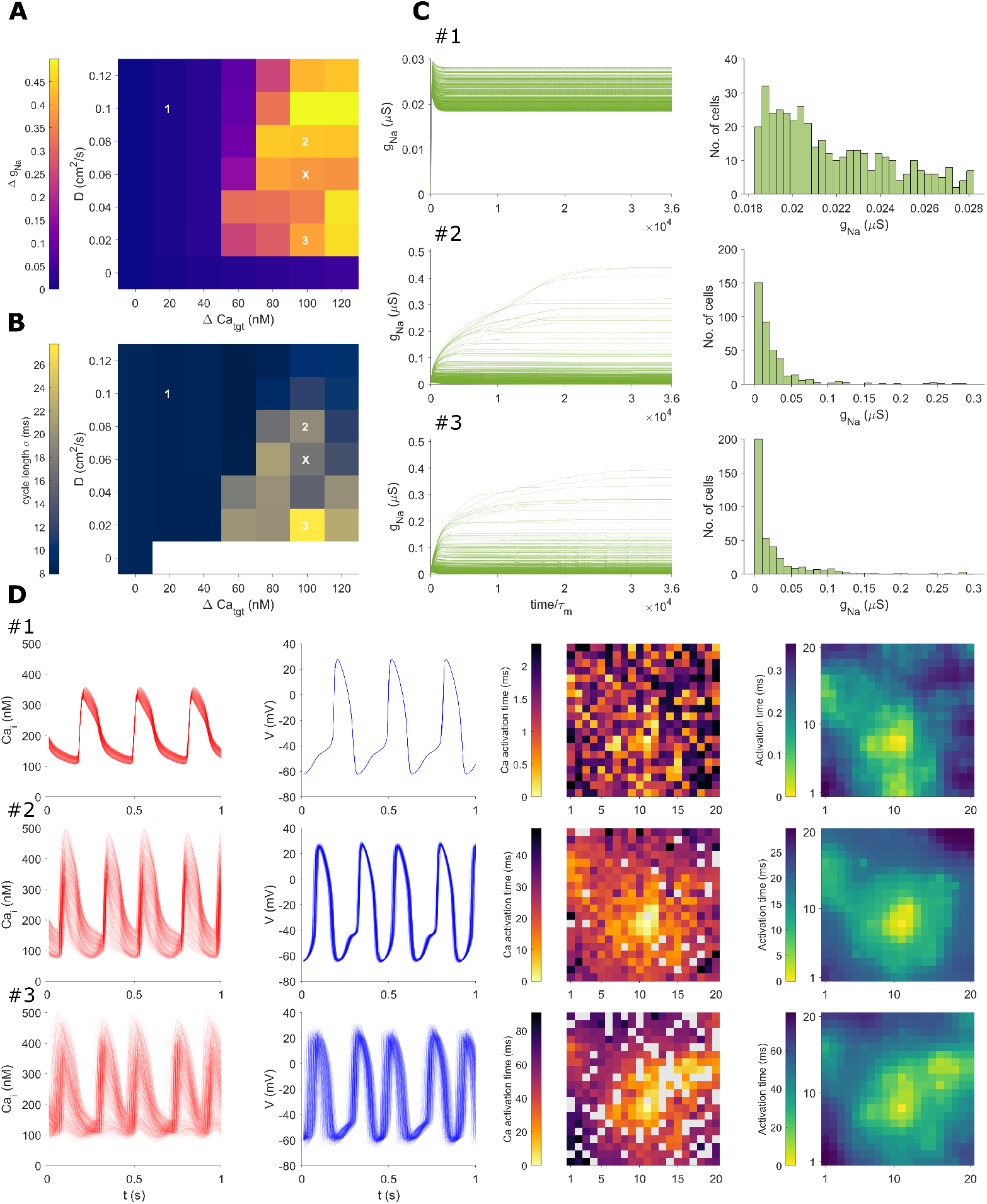
Ion channel conductance heterogeneity and cycle length variability depend on tissue electrical coupling values (*D*) and *Ca*_*tgt*_ ranges (Δ*Ca*_*tgt*_). **A**. The difference between lowest and highest *g*_*Na*_ (Δ*g*_*Na*_) and **B**. standard deviation of cycle length in the SAN tissue at steady state are shown as functions of *D* and Δ*Ca*_*tgt*_. For any *D* = 0 and any *Ca*_*tgt*_ *>* 0, all cells in tissue are completely unsynchronized, therefore there is no meaningful tissue-level cycle length (shown as white in the heatmap). **C**. Conductance time evolution and distribution and **D**. voltage and *Ca*_*i*_ activity and activation maps are shown for three different parameter pairs o1f8electrical coupling and *Ca*_*tgt*_ heterogeneity. Note the different colorbar scales in the activation maps.

As would be expected, we find that some degree of electrical coupling is necessary for synchronized electrical activity. For *D* = 0, electrical activity is equivalent to 400 individual SAN cells, and a tissue-level cycle length variability cannot be calculated (shown in white, Fig. 6B). Importantly, we also find that electrical coupling is necessary for the wide *g*_*Na*_ conductance distribution (Fig. 6A, Fig S3). That is, for *D* = 0, variation in *g*_*Na*_ follows that of individual cells, which is a much narrower distribution than that of the electrically coupled tissue. This is due the effect explained above: since diffusion couples both rhythm and AP shape, the ion channel conductances distribution across the tissue must be wide to ‘meet’ the cellular *Ca*_*tgt*_ levels.

In Fig. 6, examples 1-3 illustrate different activity patterns for different *D* or Δ*Ca*_*tgt*_ pairs (with ‘x’ denoting parameters for the conditions shown in Figs. 3-5). For strong electrical coupling and low *Ca*_*tgt*_ variability (#1, high *D*/low Δ*Ca*_*tgt*_), the resulting SAN tissue is highly homogeneous. Conductances quickly reach steady-state and remain distributed in a small range (note the small y-axis range). This leads to similar CT and voltage traces throughout the tissue, with the activation maps for calcium and voltage demonstrating almost perfect synchronization in tissue (note activation map time range) and a small cycle length variability.

For moderate electrical coupling and high *Ca*_*tgt*_ variability (#2, moderate *D*/high Δ*Ca*_*tgt*_), the SAN tissue exhibit similar behavior as the example in Figs. 3-5, with a wide conductance distribution at steady state. The CT traces are highly variable, while the voltage traces are more tightly coupled. As above, some cells exhibit negligible amplitude CT amplitudes, i.e., they do not pass the *Ca*_*i*_ threshold level (gray pixels). Finally, for weaker (but still present) electrical coupling and high *Ca*_*tgt*_ variability (#3, low *D*/high Δ*Ca*_*tgt*_), we find the largest degree of cycle length variability. This is reflected in the highly heterogeneous CTs and variable voltage traces, consistent with significantly slower activation times, compared with the prior two cases.

Overall, Figure 6 illustrate that heterogeneous channel expression and variability in cycle length emerges for a wide range of conditions, namely electrical coupling and *Ca*_*tgt*_ heterogeneity. In particular, variable cycle length requires *Ca*_*tgt*_ heterogeneity above a threshold Δ*Ca*_*tgt*_, and further requires a “sweet spot” for electrical coupling. A minimum level of electrical coupling is required for emergence of competing pacemaking clusters that drive activity across the SAN tissue, but a large degree of electrical coupling results in near perfect tissue synchronization in a manner that suppresses cycle length variability.

### SAN tissue response to perturbations and knockouts

The above results highlight that the minimal feedback model reproduces key emergent properties of spontaneous SAN tissue activity. As a final demonstration of the robustness and consistency with prior experimental studies, we consider the SAN tissue response to several types of perturbations. We first consider the SAN tissue response to injury (Fig. 7). Specifically, utilizing the SAN tissue simulation for *Ca*_*tgt*_ = 200 nM and Δ*Ca*_*tgt*_ = 100 nM (Figs. 3-5) after reaching steady-state, we mimic an ablation of the 20 cells with the highest conductance by setting the *Ca*_*tgt*_ value to 0. After a brief transient (1000 cycles) after ablation, the histogram of *g*_*Na*_ illustrates the loss of the the highest conductances (Fig. 7B). However, after the 20 ablated cells lose their function, other cells in the tissue compensate and increase their respective conductances, which leads to a new steady-state with a highly similar conductance distribution to the original one (cf. Fig. 7C and Fig. 6, #2 or Fig. 3C). After the ablation, the cycle length temporarily lengthens and loses its variability, similar to the initial period of the original simulation. However, after the compensation of the remaining cells, cycle length shortens again and becomes variable, due to new dominant pacemaking clusters in the tissue. Notably, the final average cycle length is slightly longer than the previous steady state. The voltage (Fig. 7E-F) and the CT (Fig. 7H-I) activity, as well as the final *g*_*Na*_ spatial map (Fig. 7G) are very similar to the original steady state (see Figs. 3-5). Thus, the feedback model leads to a robustness in tissue response: the cells recapitulate tissue development and qualitatively recreate the original behavior, thus exhibiting robustness to perturbations such as cell injury.

**Figure 7.**
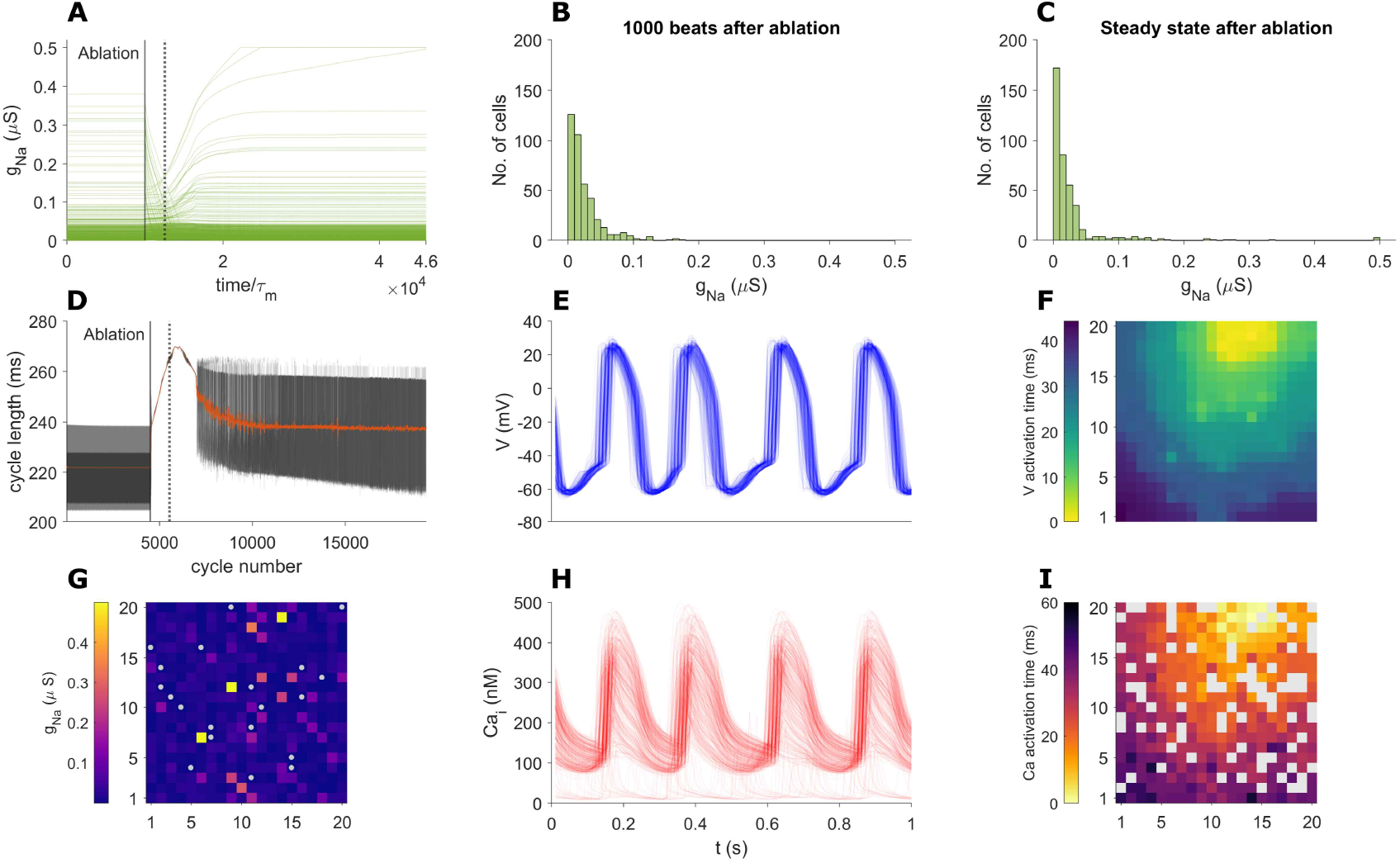
SAN tissue response to simulated pacemaking ablation. The 20 highest conductance cells are ablated, by setting *Ca*_*tgt*_ to 0, to test the robustness of the system to injury. **A**. Tissue conductance dynamics before and after ablation. **B**. Tissue conductance distribution a short time (1000 beats) after ablation (dotted vertical line in A). **C**. Tissue conductance distribution at the new steady state following ablation. **D**. Cycle length dynamics (gray) after ablation. Red line denotes a running average of the cycle length to illustrate average trends. **E**. All voltage traces at the post-ablation steady state. **F**. Example voltage activation map. **G**. Conductance spatial map at the new steady state. Ablated cells are marked by gray dots. **H**. All *Ca*_*i*_ traces at the post-ablation steady state. **I**. Example calcium activation map.

Figures 8 and 9 consider the SAN tissue response to NCX and HCN knockout, respectively (see Methods for details on KO induction). After NCX KO, all conductances decrease (Fig. 8D), driven by a transient increase in *Ca*_*i*_ (Fig. 8C). The subsequent steady-state *g*_*Na*_ distribution is similar to the original, slightly left-shifted, i.e., with overall smaller *g*_*Na*_ values (Fig. 8F). Importantly however, tissue calcium activity is significantly altered. A large number of cells fail to reach the *Ca*_*i*_ threshold (Fig. 8B), and some cells completely lose *Ca*_*i*_ oscillatory activity (Fig. 8G, #3). Other cells maintain calcium activity, but alternate a full CT with small sub-threshold peaks (Fig. 8G, #1, 2). Interestingly, voltage activity is affected to a lesser degree. The tissue average voltage amplitude decreases (Fig. 8C); however, spontaneous electrical activity still emerges from a leading pacemaking site, albeit with a longer cycle length (Fig. 8A). Importantly, the tissue loses all cycle length variability after NCX KO, as all electrical activity originates from the same pacemaking site.

**Figure 8.**
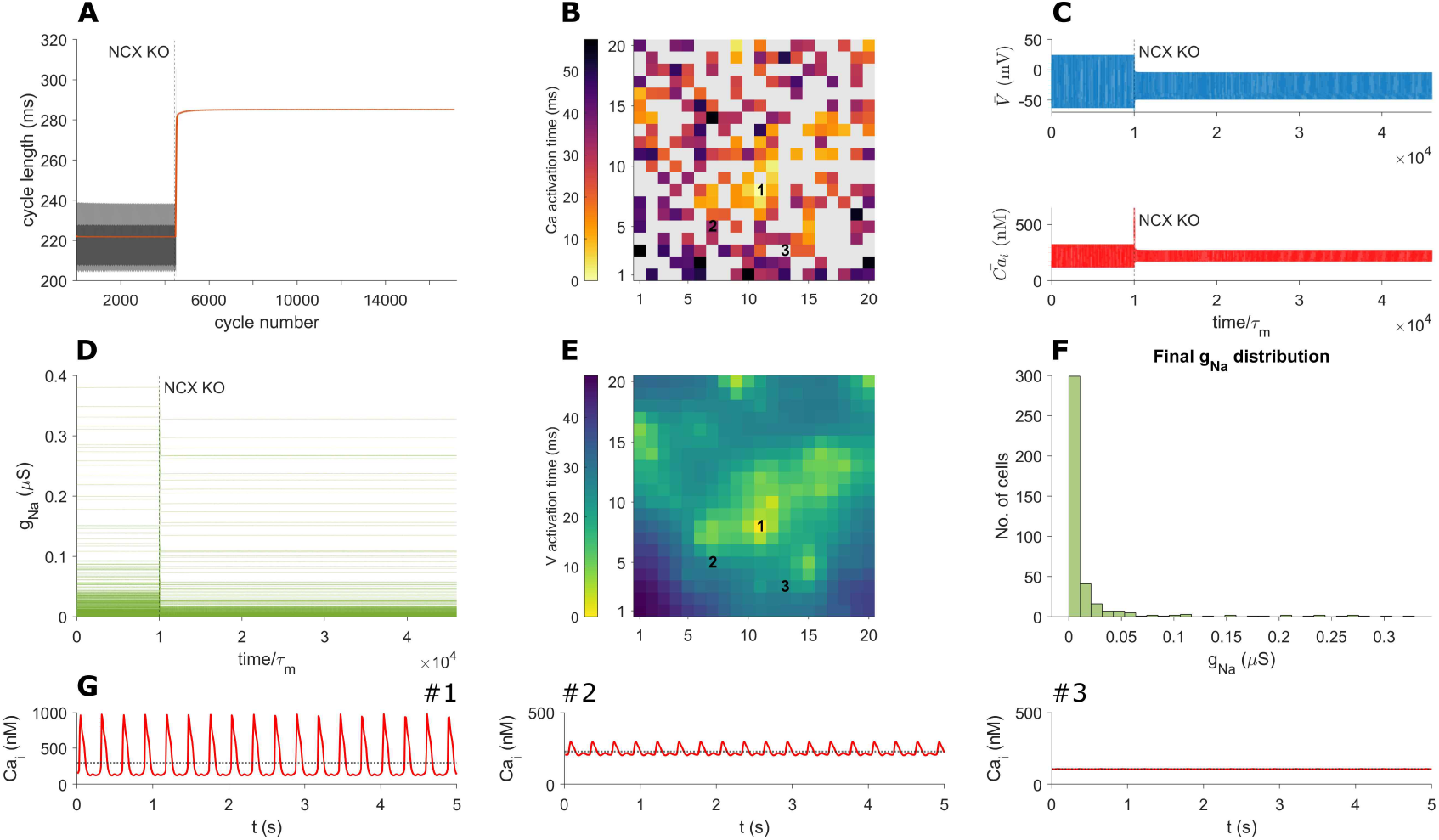
SAN tissue response following 5% NCX knockout (KO). **A**. Cycle length (gray) before and after NCX KO. Red line denotes a running average of the cycle length to illustrate average trends.**B**. *Ca*_*i*_ activation map at the new steady state. **C**. Average SAN tissue voltage and *Ca*_*i*_. **D**. Evolution of conductances before and after NCX KO. **E**. Voltage activation map at the new steady-state. **F**. Conductance distribution at the new steady-state. **G**. Example *Ca*_*i*_ traces after NCX KO at three locations, denoted in **B** and **E**. Horizontal lines denote the associated *Ca*_*tgt*_ values for each cell. Note that #3 loses all activity.

**Figure 9.**
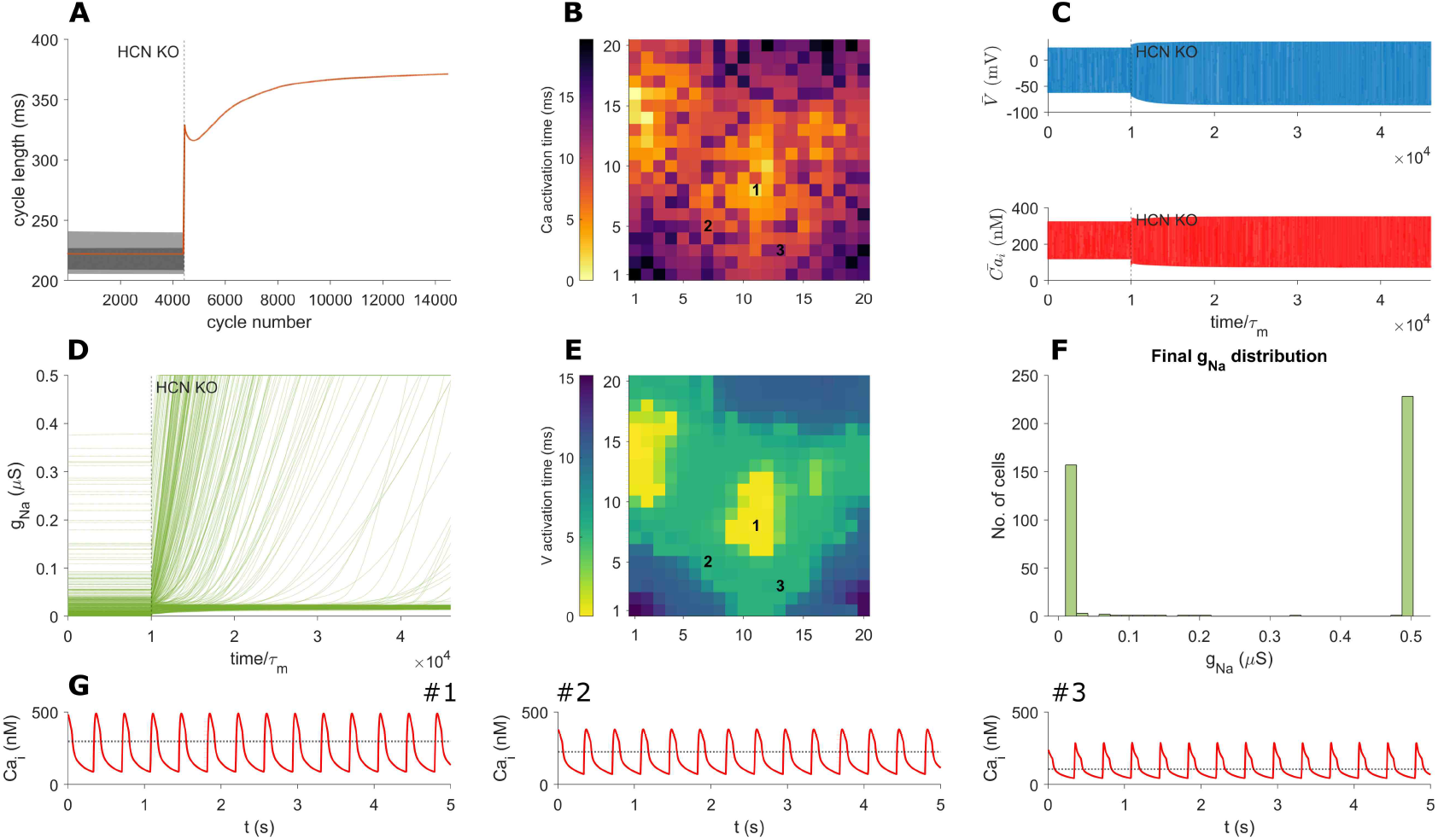
SAN tissue response to 5% HCN channel knockout (KO). **A**. Cycle length (gray) before and after NCX KO. Red line denotes a running average of the cycle length to illustrate average trends. **B**. *Ca*_*i*_ activation map at the new steady state. **C**. Average tissue voltage and *Ca*_*i*_. **D**. Evolution of conductances after HCN KO. **E**. Voltage activation map at the new steady state. **F**. New *g*_*Na*_ distribution. **G**. Example *Ca*_*i*_ traces after HCN KO at three locations, denoted in **B** and **E**. Horizontal lines denote the associated *Ca*_*tgt*_ values for each cell.

Finally, Figure 9 shows the response to HCN KO. The loss of HCN leads to a considerable increase in cell conductances (Fig. 9D), which in turn leads to increased tissue voltage and CT amplitude. This however does not translate to a shorter cycle length. On the contrary, cycle length becomes significantly longer (Fig. 9A), while, as with the NCX KO, the SAN tissue cycle length variability is also lost. Due to the increase in conductances throughout the SAN tissue, all cells demonstrate superthreshold *Ca*_*i*_ activity, and voltage activation is highly synchronized (Fig. 9B, E and G, #1-3).

## 4 Discussion

### Single SAN cell dynamics and model assumptions

In this study, we demonstrate that a calcium feedback model, coupled to a model of SAN electrical and calcium signaling, results in the emergence of spontaneous activity from an initially quiescent cell. The feedback model necessarily makes some significant simplifications of SAN biology, as similarly noted in the case of neurons by O’Leary and colleagues [30]. Here, we discuss several of these key assumptions. A first assumption is that conductance ratios are fixed between channels. This notion is in concordance with the theoretical framework of the “good enough solution”, in cardiac cells as well as neurons [29, 33]. Further, a number of studies of SAN cells have shown total conductances of key ionic currents (*I*_*f*_ [1], *I*_*CaL*_ [3] *I*_*to*_ [34], *I*_*Kr*_, *I*_*Ks*_ [35]) to be positively correlated with membrane capacitance, which is further correlated with the spontaneous rate of activity [1, 36]. This implies that larger cells have larger conductances of the mentioned currents (and thus approximately maintained proportions between them) and thus increased electrical activity. Further, larger cells also have larger calcium transients [36]. Contradicting these results, Monfredi and colleagues [4] did not find significant correlation between cell size and *I*_*f*_, *I*_*K*_ (i.e. total *K*^+^ current), *I*_*CaL*_, or cycle length. However, the authors show that the magnitude of *I*_*f*_ and *I*_*K*_ measured in the same cell are positively correlated, consistent with the ratio between these two currents maintained. Taken together, these studies strengthen the model’s assumptions of SAN channel conductance ratios. The single SAN cell results showing that conductance ratios are maintained and increased conductances lead to faster spontaneous activity (Figs. 1 and 2A, B) are therefore in good qualitative agreement with experimental data. Critically, exact ratios are not strictly necessary for the emergence of spontaneous electrical activity, highlighting the robustness of the feedback model (Fig. S2). While beyond the scope of this current study, additional regulation within the calcium feedback model could be represented by time-varying dynamics in *Ca*_*tgt*_ and/or conductance time constants *τ*_*x*_.

A second important assumption of the model is that the conductance feedback occurs on a significantly slower time scale than AP or CT dynamics. Such long-term conductance changes have been demonstrated in the context of reversible electrical remodeling in atrial fibrillation or flutter [37, 38, 39, 40] or chronic (days to weeks) rapid pacing in animal models [41, 42]. Supporting this separation of time scales, the feedback model shows that externally imposed rapid pacing leads to down-regulation of ion channels, which occurs in order for the SAN cell to maintain average *Ca*_*i*_ levels set by its inherent *Ca*_*tgt*_ value (Fig. 2D). Following the termination of external pacing, there is a transient period of increased cycle length, due to the low channel expression levels prior to conductances increasing to their previous values. This transient slow spontaneous activity is directly comparable to experiments demonstrating SAN dysfunction following rapid pacing and associated current remodeling: both *I*_*f*_ and *I*_*Ks*_ are down-regulated, alongside their respective mRNAs [41], and revert to normal levels following pacing cessation [42]. Interestingly, studies show that other channels do not exhibit changes, suggesting regulation more complex than in this model. Nonetheless, even with our simplifying assumptions, the model captures the essential qualitative dynamics of the emergence of spontaneous SAN electrical activity from quiescence, and cycle length and channel conductances changes.

### Emergent tissue heterogeneity

Extending the model to SAN tissue, individual cells exhibit a surprisingly large degree of conductance heterogeneity that naturally emerges from the feedback model (Fig. 3), more so than simply due to the heterogeneity in calcium target levels. Perhaps counterintuitively, electrical coupling leads to highly heterogeneous tissue conductances (Figs. 6 and S3). Since tissue-scale coupling inherently mitigates voltage differences between cells, thus driving spatially consistent AP morphology and frequency, the feedback model necessitates that individual cells adapt *Ca*_*i*_ levels to “match” their respective *Ca*_*tgt*_ through changes in ion channel expression. This adaptation in turn drives ion channel conductance heterogeneity and variability in CT characteristics. As shown above, this ultimately leads to multiple competing pacemaker centers, which also drive variability in tissue-level cycle length (Fig. 4). We find that it is thus essential that the SAN achieve a balance between tissue synchronization [43, 10] and low levels of electrical coupling, which are protective of the potential depolarizing influence of the surrounding atrial tissue [12, 44]. Our findings are consistent with recent computational studies similarly demonstrating that intermediate levels of electrical coupling in a heterogeneous SAN tissue promote robust automaticity [45, 46] Figure 6 shows that there is a wide band of conditions for which intermediate electrical coupling levels lead to partial synchronization in tissue and organization in distinct pacemaking centers. Notably, multiple pacemaker centers have been described experimentally in the SAN [47, 48], which can have different functions in relation to the autonomic nervous system or to the interface with the atria. The different cycle lengths appearing in the SAN tissue demonstrate a source of inherent heart rate variability of the SAN [49]. Notably, this variability and multifocal organization appears from a deterministic model; the only source of randomness in our system is the spatial patterning of the *Ca*_*tgt*_ values. The heterogeneity of SAN cells themselves has been widely reported: isolated cells have diverse current conductances (as discussed above), which lead to a wide variety of single cell cycle lengths and CTs [50, 5, 6]. Further, these studies find populations of HCN4 positive cells that are quiescent [50, 6], but can potentially be induced to beat rhythmically by β-adrenergic stimulation [50]. Isolated cells may behave quite differently than in tissue due to the loss of electrotonic interactions. As a further demonstration of this isolated cell heterogeneity, we mimic a SAN tissue cell isolation by completely decoupling cells from the end of a SAN tissue simulation, fixing the conductances at the values reached at steady-state (Fig. S4). We find significant variability in spontaneous cycle lengths, as well as in electrical and CT activity. Further, consistent with experiments, we also find a non-negligible proportion (*∼*15%) of cells with subthreshold electrical activity, which can be considered quiescent. Thus, our results show that heterogeneity in conductances and cycle lengths (including a distinct quiescent population) emerges as a consequence of the feedback model.

Finally, our simulations of heterogeneous calcium activity are particularly in good agreement with recent experiments of single cell CTs in intact SAN tissue, as described by Bychkov and colleagues [7]. The emergent activity in the SAN tissue model matches experiments in a number of key points: CTs vary widely in amplitude between cells and between beats in the same cell, with some cells almost quiescent with minimal CTs (Fig. 5, cf. Figs. 4-7 in [7]). Additionally, this diversity in activation leads to patchy, apparently discontinuous calcium activation in tissue (Fig. 5B), comparable to the spatial pattern shown in Fig. 15 from [7].

Overall, integration of the calcium feedback model to the SAN tissue explains a wide range of findings, including heterogeneity in conductances, calcium activity, and electrical activation (i.e., inherent heart rate variability and multiple pacemaking sites). Specifically, the model demonstrates that the tissue spatial patterning of the SAN into a heterogeneous network naturally appears from the self-organization of interacting cells, each of them being governed by a simple feedback mechanism. This fundamental behavior must necessarily be shaped by developmental factors [44], which create the niche in which the SAN appears and functions: the fibrous, isolated nature of the tissue, and its structural connections to the atria. Notably, our approach does not include these additional structural factors, which can further influence the exact configuration of the SAN. However, we demonstrate that the fundamental organization of the SAN cells into tissue and its persistence over time can be minimally determined by inherent cell-level regulatory mechanisms and the interactions between cells.

### Robustness to simulated injury and knockout (KO) models

Finally, we considered the effects of perturbations in SAN tissue. It is interesting to note that in the case of the ablation of the 20 highest conductance cells (Fig. 7), the feedback model not only maintained spontaneous activity, but recreates qualitatively the original conductance steady-state distribution. This further led to a comparable multifocal pattern of electrical activation, albeit with a longer average cycle length. Thus, while feedback is mediated at the cellular level, the model can still maintain tissue-level structural features.

Next, we investigated the effects of two channel knockouts, NCX and HCN. There are a number of studies that describe NCX KO in the mouse: Hermann and colleagues showed that an inducible NCX KO leads to progressive bradycardia, which stabilizes after 4 weeks [51], while isolated cells show arrhythmic oscillations and lower amplitude CTs. Groenke and colleagues described an atrial-specific KO in which P-waves are completely abolished [52]. Isolated SAN cells were responsive to pacing and had shorter AP duration, but unchanged CTs. Importantly, *I*_*f*_ was similar to control, but *I*_*CaL*_ was reduced. Using the same mouse model, Torrente and colleagues imaged the intact SAN, showing bursting electrical and calcium oscillations [53]. However, the depolarization waves cannot escape to the atria, likely due to significant fibrosis of both atria and SAN. Between bursts, cells also exhibited significant sub-cellular calcium releases.

The feedback model of the SAN tissue with NCX KO shows similarities to these experimental results (Fig. 8). After inducing KO, there is significant bradycardia, after a short period of adaptation. All cell conductances drop slightly, leading to significantly smaller APs and CTs. However, Ca^2+^ homeostasis is generally maintained, especially compared to simulations of NCX KO without the feedback model (Figs. S5 and S8), which show greatly increased [*Ca*]_*i*_. Moreover, in some cells (Fig. 8G #1, 2), there are small Ca^2+^ peaks between transients. Further, in single cell simulations (Fig. S7C) and in tissue (Fig. 8G #3) completely quiescent cells can appear. Finally, progressively increased KO severity leads to complete loss of both electrical and calcium activity, even with conductance compensation (Fig. S7).

HCN KOs have also been previously studied in mice. Hermann [54] and Hoesl [55] investigated a partial KO, with approximately 25% of *I*_*f*_ remaining after induction. Both studies report overall maintained heart rates, but frequent sinus pauses. Baruscotti and colleagues show that, on the contrary, similar levels of HCN KO leads to progressive bradycardia followed by the death of the animals [56]. The SAN activity is significantly bradycardic; however the ultimate cause of heart arrest seems to be atrioventricular node block. Mesirca and colleagues reported that a complete abolition of *I*_*f*_, through overexpression of a non-functional HCN channel, led to significantly reduced isolated cell activity, accompanied by frequent pauses [57]. Moreover, there are Ca^2+^ abnormalities: *I*_*CaL*_ is increased, and CTs have larger amplitudes. The mice are also significantly bradycardic, although the difference between wildtype and KOs is attenuated by complete autonomic block.

The feedback model predicts that HCN KO leads to bradycardia in tissue (Fig. 9). The cells compensate by significantly increasing channel conductances, up to an upper threshold imposed. However, even after channel up-regulation, the model is not able to compensate for the lack of *I*_*f*_ and restore normal cycle length. The conductance increase also leads to significantly higher amplitude CTs even in formerly quiescent cells. Inducing KO without feedback compensation (Figs S6 and S10) also leads to bradycardia, but with drastically perturbed Ca^2+^ homeostasis. Interestingly, in single cells, a partial KO of 20% (Fig. S11) leads to very similar results as the ones described by Hermann and Hoesl cited above: The average cycle length is not significantly changed, but the cell exhibits variable cycle lengths with pauses between beats. Without the feedback model, the cell is significantly bradycardic and loses Ca^2+^ homeostasis. Further, as KO severity increases (Fig. S9), the cells become progressively more bradycardic, as the feedback model loses its ability to sufficiently compensate for the loss of *I*_*f*_ .

There are however important limitations of the feedback model. It does not recreate the bursting behavior of NCX KO tissue, or the almost perfect compensation in complete HCN KOs. Further, some studies show no gene expression changes after KO [57], or minimal current changes [52] that the feedback model predicts would be significantly altered. Even with these limitations, the model still captures many key aspects of adaptation after induced KOs. Significantly, the feedback model succeeds in capturing some key qualitative results, as well as maintaining Ca^2+^ activity within normal bounds in both KO cases. Further, even though both KOs lead to bradycardia, experiments demonstrate opposite directional changes in *I*_*CaL*_ (decrease in the NCX KO and increase in the HCN KO), both of which are accurately predicted by the feedback model.

We note that there are other adaptations that are not included in our study, such as tissue-level fibrosis [53] or interactions with the autonomic nervous system [52], which can significantly modulate the behavior of the SAN. Finally, all the KO studies have been carried out in mice, while here we simulate the rabbit SAN cell, leading to potentially important unaccounted species differences. Despite these limitations, the feedback model reproduces many key aspects of experimentally-demonstrated perturbations to the SAN tissue.

## Conclusion and future directions

In this study, we adapted for the first time a dynamical feedback model to the SAN cell, bridging the short term AP time scale with the long term regulation of ion channel expression and conductances. Starting from simple assumptions of the underlying regulatory processes, we show that the feedback model recreates key aspects of SAN structure and function: consistent spontaneous oscillatory activity; the emergence and long-term stability of significant tissue heterogeneity and multiple pacemaking centers; the generation of coherent but inherently variable electrical activation from this heterogeneous structure; and the robustness of the system to cell injury or channel KOs. It is significant that both tissue-level structure and activity can emerge from simple cell-level feedback regulation. Further, the tissue can maintain homeostasis following perturbations just from self-organizing interactions between cells.

Precisely because the assumptions are minimal, the feedback model can be built upon and refined, including further components such as detailed intracellular regulatory networks, higher-level factors such as SAN structure, its relation to the atria, or interactions with the nervous system. The feedback model thus provides a basic framework in which to study long term structural and electrical development and remodeling in the SAN.

## Author contributions

NM and SHW designed the research. NM performed the research. NM wrote the original draft of the manuscript. SHW revised the final manuscript. All authors have read and approved the final version of this manuscript and agree to be accountable for all aspects of the work in ensuring that questions related to the accuracy or the integrity of any part of the work were properly investigated and resolved. All persons designated as authors qualify for authorship, and all those who qualify for authorship are listed.

## Acknowledgements

None.

## References

[1] H Honjo, M R Boyett, I Kodama, and J Toyama. Correlation between electrical activity and the size of rabbit sino-atrial node cells. The Journal of Physiology, 496(3):795–808, November 1996.

[2] M Boyett. The sinoatrial node, a heterogeneous pacemaker structure. Cardiovascular Research, 47(4):658–687, September 2000.

[3] Hanny Musa, Ming Lei, Hauro Honjo, Sandra A. Jones, Halina Dobrzynski, Mathew K. Lancaster, Yoshiko Takagishi, Zaineb Henderson, Itsuo Kodama, and Mark R. Boyett. Heterogeneous Expression of Ca 2+ Handling Proteins in Rabbit Sinoatrial Node. Journal of Histochemistry & Cytochemistry, 50(3):311–324, March 2002.

[4] Oliver Monfredi, Kenta Tsutsui, Bruce Ziman, Michael D. Stern, Edward G. Lakatta, and Victor A. Maltsev. Electrophysiological heterogeneity of pacemaker cells in the rabbit intercaval region, including the SA node: Insights from recording multiple ion currents in each cell. American Journal of Physiology-Heart and Circulatory Physiology, 314(3):H403–H414, March 2018.

[5] Mary S. Kim, Alexander V. Maltsev, Oliver Monfredi, Larissa A. Maltseva, Ashley Wirth, Maria Cristina Florio, Kenta Tsutsui, Daniel R. Riordon, Sean P. Parsons, Syevda Tagirova, Bruce D. Ziman, Michael D. Stern, Edward G. Lakatta, and Victor A. Maltsev. Heterogeneity of calcium clock functions in dormant, dysrhythmically and rhythmically firing single pacemaker cells isolated from SA node. Cell Calcium, 74:168–179, September 2018.

[6] Nathan Grainger, Laura Guarina, Robert H Cudmore, and L Fernando Santana. The Organization of the Sinoatrial Node Microvasculature Varies Regionally to Match Local Myocyte Excitability. Function, 2(4):zqab031, July 2021.

[7] Rostislav Bychkov, Magdalena Juhaszova, Kenta Tsutsui, Christopher Coletta, Michael D. Stern, Victor A. Maltsev, and Edward G. Lakatta. Synchronized Cardiac Impulses Emerge From Heterogeneous Local Calcium Signals Within and Among Cells of Pacemaker Tissue. JACC: Clinical Electrophysiology, 6(8):907–931, August 2020.

[8] H. Dobrzynski, J. Li, J. Tellez, I.D. Greener, V.P. Nikolski, S.E. Wright, S.H. Parson, S.A. Jones, M.K. Lancaster, M. Yamamoto, H. Honjo, Y. Takagishi, I. Kodama, I.R. Efimov, R. Billeter, and M.R. Boyett. Computer Three-Dimensional Reconstruction of the Sinoatrial Node. Circulation, 111(7):846–854, February 2005.

[9] M.R. Boyett, S. Inada, S. Yoo, J. Li, J. Liu, J. Tellez, I.D. Greener, H. Honjo, R. Billeter, M. Lei, H. Zhang, I.R. Efimov, and H. Dobrzynski. Connexins in the Sinoatrial and Atrioventricular Nodes. In S. Dhein, editor, Advances in Cardiology, pages 175–197. KARGER, Basel, 2006.

[10] D C Michaels, E P Matyas, and J Jalife. Mechanisms of sinoatrial pacemaker synchronization: A new hypothesis. Circulation Research, 61(5):704–714, November 1987.

[11] E. Etienne Verheijck, Ronald Wilders, Ronald W. Joyner, David A. Golod, Rajiv Kumar, Habo J. Jongsma, Lennart N. Bouman, and Antoni C.G. van Ginneken. Pace-maker Synchronization of Electrically Coupled Rabbit Sinoatrial Node Cells. Journal of General Physiology, 111(1):95–112, January 1998.

[12] Sathya D. Unudurthi, Roseanne M. Wolf, and Thomas J. Hund. Role of sinoatrial node architecture in maintaining a balanced source-sink relationship and synchronous cardiac pacemaking. Frontiers in Physiology, 5, November 2014.

[13] Amrita X. Sarkar, David J. Christini, and Eric A. Sobie. Exploiting mathematical models to illuminate electrophysiological variability between individuals: Electrophysiological variability. The Journal of Physiology, 590(11):2555–2567, June 2012.

[14] Astrid A Prinz, Dirk Bucher, and Eve Marder. Similar network activity from disparate circuit parameters. Nature Neuroscience, 7(12):1345–1352, December 2004.

[15] Amrita X. Sarkar and Eric A. Sobie. Regression Analysis for Constraining Free Parameters in Electrophysiological Models of Cardiac Cells. PLoS Computational Biology, 6(9):e1000914, September 2010.

[16] James N. Weiss, Alain Karma, W. Robb MacLellan, Mario Deng, Christoph D. Rau, Colin M. Rees, Jessica Wang, Nicholas Wisniewski, Eleazar Eskin, Steve Horvath, Zhilin Qu, Yibin Wang, and Aldons J. Lusis. “Good Enough Solutions” and the Genetics of Complex Diseases. Circulation Research, 111(4):493–504, August 2012.

[17] Gwendal LeMasson, Eve Marder, and L. F. Abbott. Activity-Dependent Regulation of Conductances in Model Neurons. Science, 259(5103):1915–1917, March 1993.

[18] C. Gunay and A. A. Prinz. Model Calcium Sensors for Network Homeostasis: Sensor and Readout Parameter Analysis from a Database of Model Neuronal Networks. Journal of Neuroscience, 30(5):1686–1698, February 2010.

[19] Andrey V. Olypher and Astrid A. Prinz. Geometry and dynamics of activity-dependent homeostatic regulation in neurons. Journal of Computational Neuroscience, 28(3):361–374, June 2010.

[20] Victor A. Maltsev and Edward G. Lakatta. Synergism of coupled subsarcolemmal Ca 2+ clocks and sarcolemmal voltage clocks confers robust and flexible pacemaker function in a novel pacemaker cell model. American Journal of Physiology-Heart and Circulatory Physiology, 296(3):H594–H615, March 2009.

[21] Anna V. Maltsev, Victor A. Maltsev, Maxim Mikheev, Larissa A. Maltseva, Syevda G. Sirenko, Edward G. Lakatta, and Michael D. Stern. Synchronization of Stochastic Ca2+ Release Units Creates a Rhythmic Ca2+ Clock in Cardiac Pacemaker Cells. Biophysical Journal, 100(2):271–283, January 2011.

[22] Damian G. Wheeler, Rachel D. Groth, Huan Ma, Curtis F. Barrett, Scott F. Owen, Parsa Safa, and Richard W. Tsien. CaV1 and CaV2 Channels Engage Distinct Modes of Ca2+ Signaling to Control CREB-Dependent Gene Expression. Cell, 149(5):1112–1124, May 2012.

[23] Timothy O’Leary, Mark C. W. van Rossum, and David J. A. Wyllie. Homeostasis of intrinsic excitability in hippocampal neurones: Dynamics and mechanism of the response to chronic depolarization: Homeostatic regulation of intrinsic excitability. The Journal of Physiology, 588(1):157–170, January 2010.

[24] Barbara Rosati and David McKinnon. Regulation of Ion Channel Expression. Circulation Research, 94(7):874–883, April 2004.

[25] Xiao Yan Qi, Yung-Hsin Yeh, Ling Xiao, Brett Burstein, Ange Maguy, Denis Chartier, Louis R. Villeneuve, Bianca J.J.M. Brundel, Dobromir Dobrev, and Stanley Nattel. Cel-lular Signaling Underlying Atrial Tachycardia Remodeling of L-type Calcium Current. Circulation Research, 103(8):845–854, October 2008.

[26] Samantha C. Salvage, Zaki F. Habib, Hugh R. Matthews, Antony P. Jackson, and Christopher L.-H. Huang. Ca2+-dependent modulation of voltage-gated myocyte sodium channels. Biochemical Society Transactions, 49(5):1941–1961, November 2021.

[27] Yuejin Wu and Mark E. Anderson. CaMKII in sinoatrial node physiology and dysfunction. Frontiers in Pharmacology, 5, March 2014.

[28] Jie Liu, Syevda Sirenko, Magdalena Juhaszova, Bruce Ziman, Veena Shetty, Silvia Rain, Shweta Shukla, Harold A. Spurgeon, Tatiana M. Vinogradova, Victor A. Maltsev, and Edward G. Lakatta. A full range of mouse sinoatrial node AP firing rates requires protein kinase A-dependent calcium signaling. Journal of Molecular and Cellular Cardiology, 51(5):730–739, November 2011.

[29] Colin M Rees, Jun-Hai Yang, Marc Santolini, Aldons J Lusis, James N Weiss, and Alain Karma. The Ca2+ transient as a feedback sensor controlling cardiomyocyte ionic conductances in mouse populations. eLife, 7:e36717, September 2018.

[30] Timothy O’Leary, Alex H. Williams, Alessio Franci, and Eve Marder. Cell Types, Network Homeostasis, and Pathological Compensation from a Biologically Plausible Ion Channel Expression Model. Neuron, 82(4):809–821, May 2014.

[31] Stefano Severi, Matteo Fantini, Lara A. Charawi, and Dario DiFrancesco. An updated computational model of rabbit sinoatrial action potential to investigate the mechanisms of heart rate modulation: Model of SAN action potential. The Journal of Physiology, 590(18):4483–4499, September 2012.

[32] Victor A. Maltsev, Tatiana M. Vinogradova, and Edward G. Lakatta. The Emergence of a General Theory of the Initiation and Strength of the Heartbeat. Journal of Pharmacological Sciences, 100(5):338–369, 2006.

[33] T. O’Leary, A. H. Williams, J. S. Caplan, and E. Marder. Correlations in ion channel expression emerge from homeostatic tuning rules. Proceedings of the National Academy of Sciences, 110(28):E2645–E2654, July 2013.

[34] M Lei. Characterisation of the transient outward K+ current in rabbit sinoatrial node cells. Cardiovascular Research, 46(3):433–441, June 2000.

[35] M. Lei, H. Honjo, I. Kodama, and M. R. Boyett. Heterogeneous expression of the delayed-rectifier K + currents i K,r and i K,s in rabbit sinoatrial node cells. The Journal of Physiology, 535(3):703–714, September 2001.

[36] Matthew K. Lancaster, Sandra A. Jones, Simon M. Harrison, and Mark R. Boyett. In-tracellular Ca 2+ and pacemaking within the rabbit sinoatrial node: Heterogeneity of role and control: Heterogeneity of [Ca 2+] i within the SAN. The Journal of Physiology, 556(2):481–494, April 2004.

[37] Arif Elvan, Kevin Wylie, and Douglas P. Zipes. Pacing-Induced Chronic Atrial Fibrillation Impairs Sinus Node Function in Dogs: Electrophysiological Remodeling. Circulation, 94(11):2953–2960, December 1996.

[38] Stanley Nattel, Ange Maguy, Sabrina Le Bouter, and Yung-Hsin Yeh. Arrhythmogenic Ion-Channel Remodeling in the Heart: Heart Failure, Myocardial Infarction, and Atrial Fibrillation. Physiological Reviews, 87(2):425–456, April 2007.

[39] Stanley Nattel and Masahide Harada. Atrial Remodeling and Atrial Fibrillation. Journal of the American College of Cardiology, 63(22):2335–2345, June 2014.

[40] Michael R Franz, Pamela L Karasik, Cuilan Li, Jean Moubarak, and Mary Chavez. Electrical Remodeling of the Human Atrium: Similar Effects in Patients With Chronic Atrial Fibrillation and Atrial Flutter. Journal of the American College of Cardiology, 30(7):1785–1792, December 1997.

[41] Yung-Hsin Yeh, Brett Burstein, Xiao Yan Qi, Masao Sakabe, Denis Chartier, Philippe Comtois, Zhiguo Wang, Chi-Tai Kuo, and Stanley Nattel. Funny Current Downregulation and Sinus Node Dysfunction Associated With Atrial Tachyarrhythmia: A Molecular Basis for Tachycardia-Bradycardia Syndrome. Circulation, 119(12):1576–1585, March 2009.

[42] Zhisong Chen, Bing Sun, Gary Tse, Jinfa Jiang, and Wenjun Xu. Reversibility of both sinus node dysfunction and reduced HCN4 mRNA expression level in an atrial tachycardia pacing model of tachycardia-bradycardia syndrome in rabbit hearts. International Journal of Clinical and Experimental Pathology, page 6, 2015.

[43] Daniel Gratz, Birce Onal, Alyssa Dalic, and Thomas J. Hund. Synchronization of Pacemaking in the Sinoatrial Node: A Mathematical Modeling Study. Frontiers in Physics, 6:63, June 2018.

[44] Marietta Easterling, Simone Rossi, Anthony J Mazzella, and Michael Bressan. Assembly of the Cardiac Pacemaking Complex: Electrogenic Principles of Sinoatrial Node Morphogenesis. Journal of Cardiovascular Development and Disease, 8(4):40, April 2021.

[45] Chiara Campana, Eugenio Ricci, Chiara Bartolucci, Stefano Severi, and Eric A. Sobie. Coupling and Heterogeneity Modulate Pacemaking Capability in Healthy and Diseased Two-Dimensional Sinoatrial Node Tissue Models. Preprint, Physiology, April 2022.

[46] Alexander V Maltsev, Michael D Stern, Edward G Lakatta, and Victor A Maltsev. Functional heterogeneity of cell populations increases robustness of pacemaker function in a numerical model of the sinoatrial node tissue. Frontiers in physiology, page 605, 2022.

[47] Vadim V. Fedorov, Alexey V. Glukhov, and Roger Chang. Conduction barriers and pathways of the sinoatrial pacemaker complex: Their role in normal rhythm and atrial arrhythmias. American Journal of Physiology-Heart and Circulatory Physiology, 302(9):H1773–H1783, May 2012.

[48] Jaclyn A. Brennan, Qing Chen, Anna Gams, Jhansi Dyavanapalli, David Mendelowitz, Weiqun Peng, and Igor R. Efimov. Evidence of Superior and Inferior Sinoatrial Nodes in the Mammalian Heart. JACC: Clinical Electrophysiology, 6(14):1827–1840, December 2020.

[49] Aviv A. Rosenberg, Ido Weiser-Bitoun, George E. Billman, and Yael Yaniv. Signatures of the autonomic nervous system and the heart’s pacemaker cells in canine electrocardiograms and their applications to humans. Scientific Reports, 10(1):9971, December 2020.

[50] Mary S. Kim, Oliver Monfredi, Larissa A. Maltseva, Edward G. Lakatta, and Victor A. Maltsev. β-Adrenergic Stimulation Synchronizes a Broad Spectrum of Action Potential Firing Rates of Cardiac Pacemaker Cells toward a Higher Population Average. Cells, 10(8):2124, August 2021.

[51] Stefan Herrmann, Peter Lipp, Kathrina Wiesen, Juliane Stieber, Huong Nguyen, Elisabeth Kaiser, and Andreas Ludwig. The cardiac sodium–calcium exchanger NCX1 is a key player in the initiation and maintenance of a stable heart rhythm. Cardiovascular Research, 99(4):780–788, September 2013.

[52] Sabine Groenke, Eric D. Larson, Sarah Alber, Rui Zhang, Scott T. Lamp, Xiaoyan Ren, Haruko Nakano, Maria C. Jordan, Hrayr S. Karagueuzian, Kenneth P. Roos, Atsushi Nakano, Catherine Proenza, Kenneth D. Philipson, and Joshua I. Goldhaber. Complete Atrial-Specific Knockout of Sodium-Calcium Exchange Eliminates Sinoatrial Node Pacemaker Activity. PLoS ONE, 8(11):e81633, November 2013.

[53] Angelo G. Torrente, Rui Zhang, Audrey Zaini, Jorge F. Giani, Jeanney Kang, Scott T. Lamp, Kenneth D. Philipson, and Joshua I. Goldhaber. Burst pacemaker activity of the sinoatrial node in sodium–calcium exchanger knockout mice. Proceedings of the National Academy of Sciences, 112(31):9769–9774, August 2015.

[54] Stefan Herrmann, Juliane Stieber, Georg Stöckl, Franz Hofmann, and Andreas Lud-wig. HCN4 provides a ‘depolarization reserve’ and is not required for heart rate acceleration in mice. The EMBO Journal, 26(21):4423–4432, October 2007.

[55] Evelyn Hoesl, Juliane Stieber, Stefan Herrmann, Susanne Feil, Elisabeth Tybl, Franz Hofmann, Robert Feil, and Andreas Ludwig. Tamoxifen-inducible gene deletion in the cardiac conduction system. Journal of Molecular and Cellular Cardiology, 45(1):62–69, July 2008.

[56] M. Baruscotti, A. Bucchi, C. Viscomi, G. Mandelli, G. Consalez, T. Gnecchi-Rusconi, N. Montano, K. R. Casali, S. Micheloni, A. Barbuti, and D. DiFrancesco. Deep bradycardia and heart block caused by inducible cardiac-specific knockout of the pacemaker channel gene Hcn4. Proceedings of the National Academy of Sciences, 108(4):1705–1710, January 2011.

[57] Pietro Mesirca, Jacqueline Alig, Angelo G. Torrente, Jana Christina Müller, Laurine Marger, Anne Rollin, Claire Marquilly, Anne Vincent, Stefan Dubel, Isabelle Bidaud, Anne Fernandez, Anika Seniuk, Birgit Engeland, Jasmin Singh, Lucile Miquerol, Heimo Ehmke, Thomas Eschenhagen, Joel Nargeot, Kevin Wickman, Dirk Isbrandt, and Matteo E. Mangoni. Cardiac arrhythmia induced by genetic silencing of ‘funny’ (f) channels is rescued by GIRK4 inactivation. Nature Communications, 5(1):4664, December 2014.

