## Supplemental Material for "Emergent Electrical Activity, Tissue Heterogeneity, and Robustness in a Calcium Feedback Regulatory Model of the Sinoatrial Node"

### Supplemental Figures

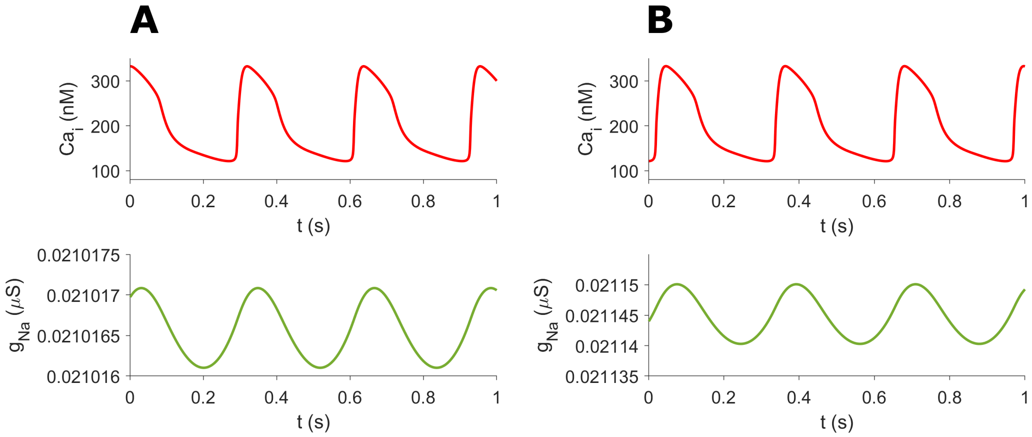


**Figure S1.** Changes in ion conductances during calcium transient (CT) oscillations. Top: CTs at steady state. Bottom: variation of $g_{Na}$ conductance with (**A**) single cell and (**B**) tissue $\tau_{g_{Na}}$ values at steady state. Note the slightly different y-axis values and scale.

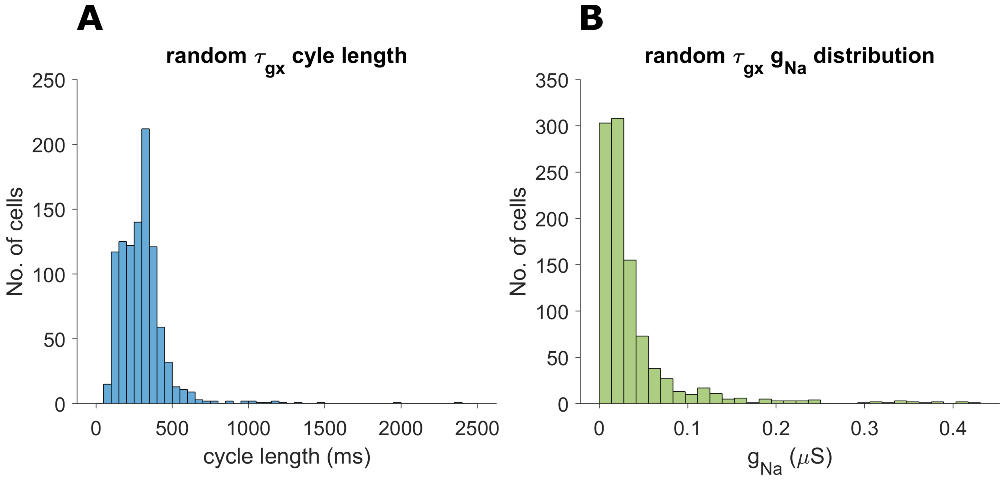


**Figure S2.** Histograms of spontaneous activity and conductance distributions from 1000 single cell simulations with random time constant $\tau_{gx}$. **A.** Distribution of spontaneous cycle lengths, from the 998 out of 1000 cells that exhibit oscillatory activity. **B.** Steady-state $g_{Na}$ distribution.

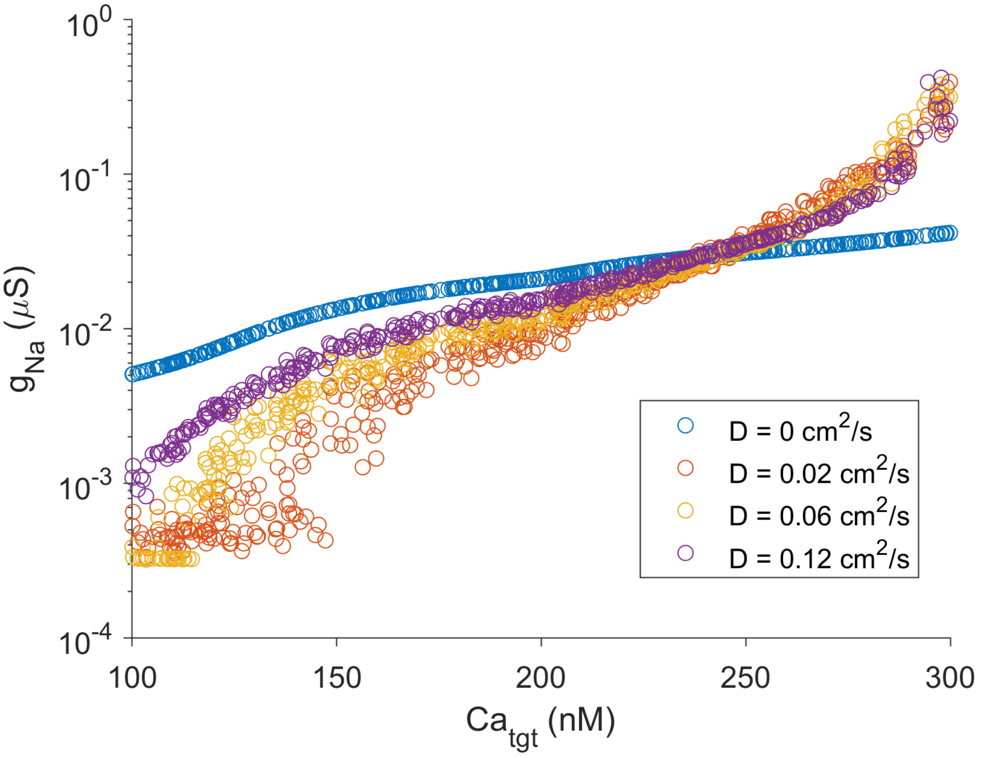


**Figure S3.** Scatter plot showing the relationship between the cell $Ca_{tgt}$ and $g_{Na}$ at steady state, for different $D$ tissue coupling values. $D$ above 0 greatly increases heterogeneity compared with single cells (i.e., $D = 0$); however further increasing $D$ moderately reduces tissue heterogeneity. Note the logarithmic scale for the y-axis.

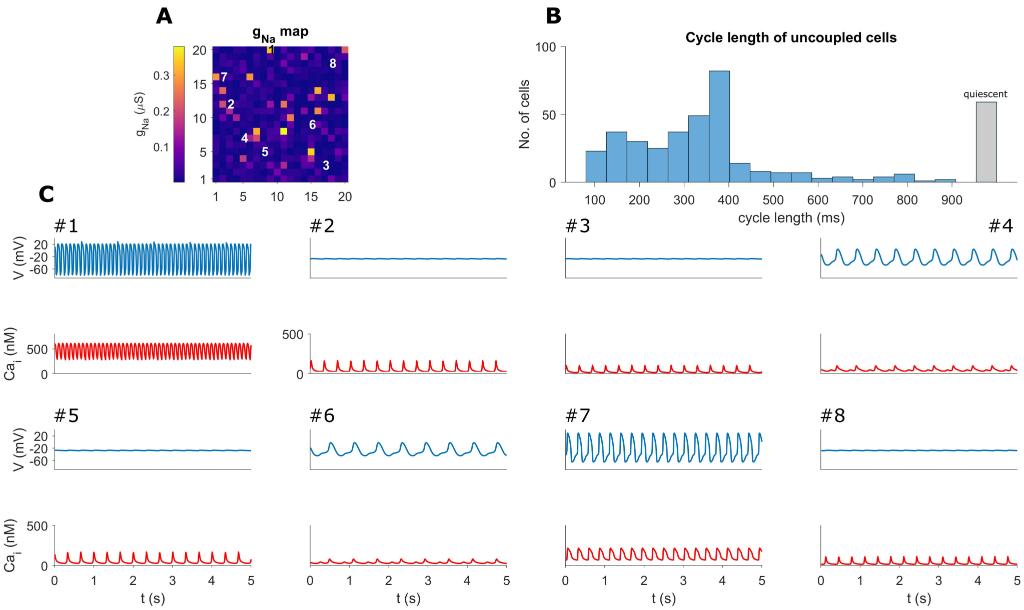


**Figure S4.** Simulations showing the behavior of isolated cells from tissue. Cell isolation is simulated by setting $D = 0$ and fixing all conductances at steady state values. **A.** $g_{Na}$ map for reference, showing the fixed steady state values. **B.** Cycle length distribution of isolated cells. 59 cells out of 400 (14.75%) have no oscillatory activity (grey bar). **C.** Example voltage (blue) and calcium (red) traces of isolated cells. Loss of tissue coupling allows the AP shape and frequency to vary. Cells #2, 3, 5, 8 show no significant oscillations. Cells #4, 6 show subthreshold $Ca_i$ activity between complete APs. Cells #2, 3, 5, 8 show calcium oscillations, even though they lacks APs. Note that the example traces are the same as in Figure 5.

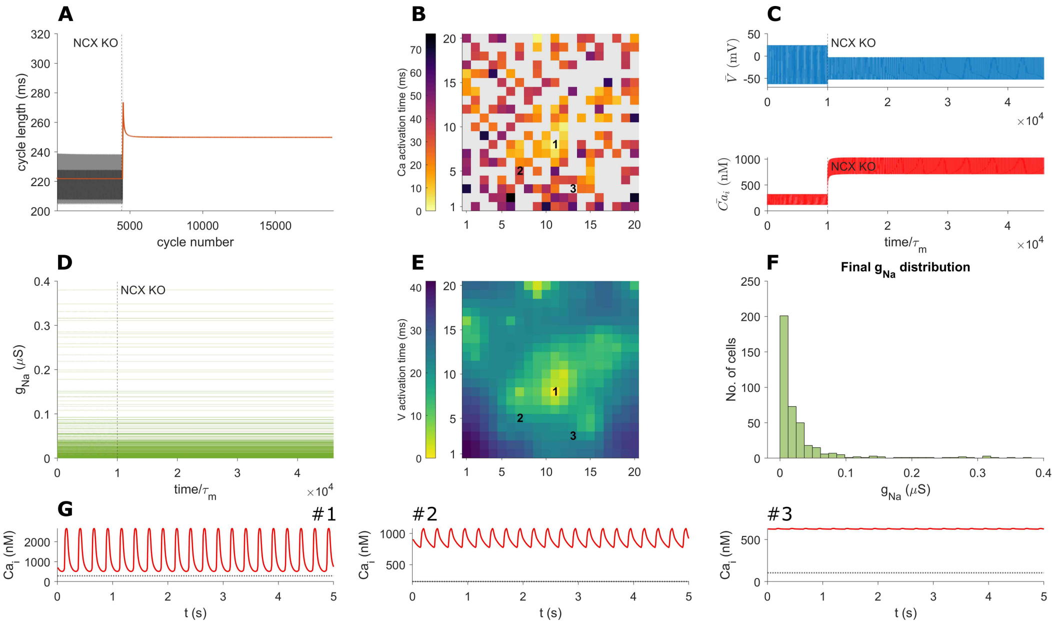


**Figure S5.** Tissue NCX KO without feedback (conductances fixed at steady state). **A.** Cycle length (gray) before and after NCX KO. Red line denotes a running average of the cycle length to illustrate average trends.**B.** $Ca_i$ activation map at the new steady state. **C.** Average SAN tissue voltage and $Ca_i$. **D.** Conductances are fixed at steady state after KO. **E.** Voltage activation map. **F.** Conductance distribution at steady-state. **G.** Example $Ca_i$ traces after NCX KO with fixed conductances at three locations, denoted in **B** and **E**. Note that #3 loses all activity. Horizontal lines denote the associated $Ca_{tgt}$ values for each cell. Note that since conductances are fixed, the CTs are no longer centered on the $Ca_{tgt}$

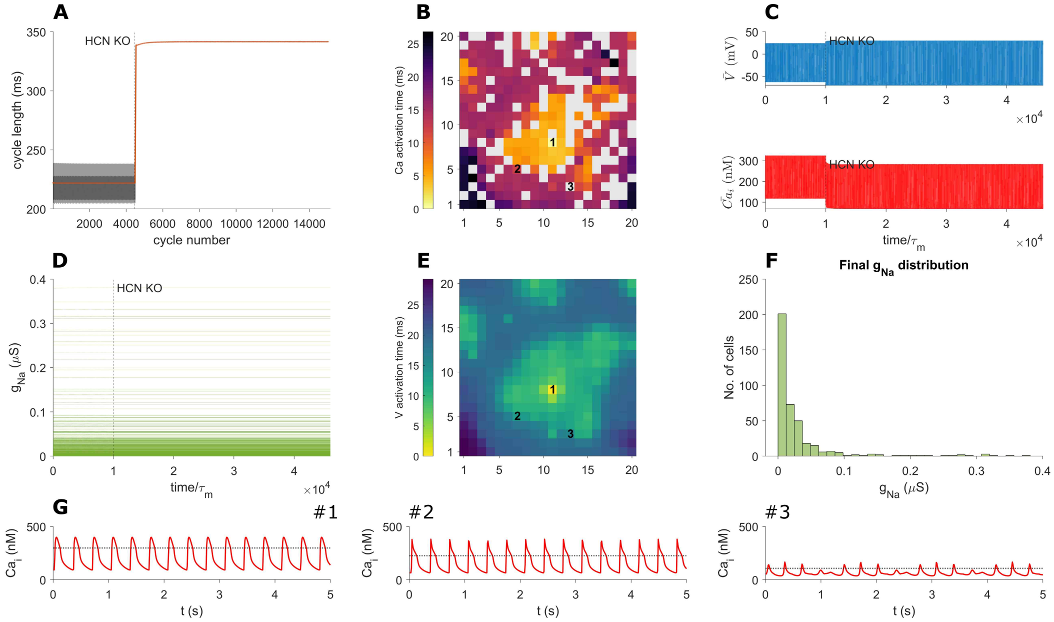


**Figure S6.** Tissue HCN KO without feedback (conductances fixed at steady state). **A.** Cycle length (gray) before and after HCN KO. Red line denotes a running average of the cycle length to illustrate average trends. **B.** $Ca_i$ activation map after KO **C.** Average tissue voltage and $Ca_i$. **D.** Conductances are fixed at steady state after KO. **E.** Voltage activation map **F.** Conductance distribution at steady-state. **G.** Example $Ca_i$ traces after HCN KO at three locations, denoted in **B** and **E**. Horizontal lines denote the associated $Ca_{tgt}$ values for each cell. Note that since conductances are fixed, the CTs are no longer centered on the $Ca_{tgt}$

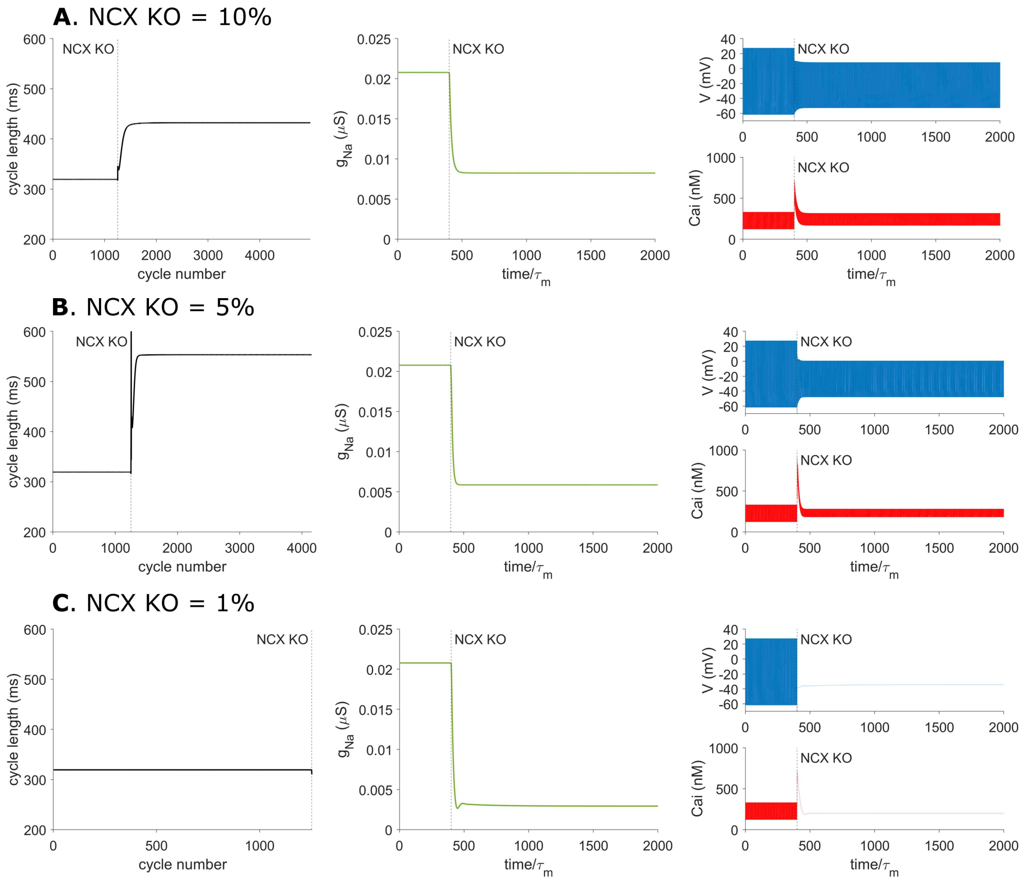


**Figure S7.** NCX KO in the single cell, for three different KO levels. Panels show from left to right: cell cycle length, $g_{Na}$, cell $V$ and $Ca_i$.

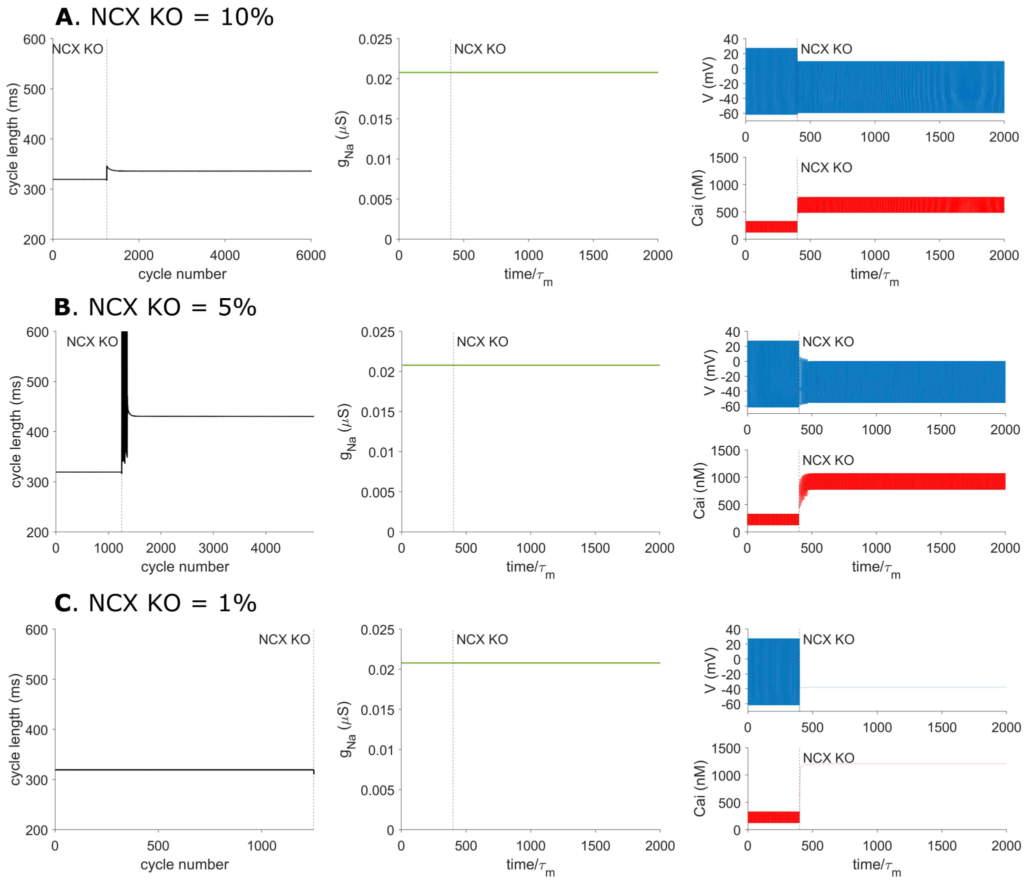


**Figure S8.** NCX KO in the single cell without feedback (fixed steady state conductances), for three different KO levels. Panels show from left to right: cell cycle length, $g_{Na}$, cell $V$ and $Ca_i$.

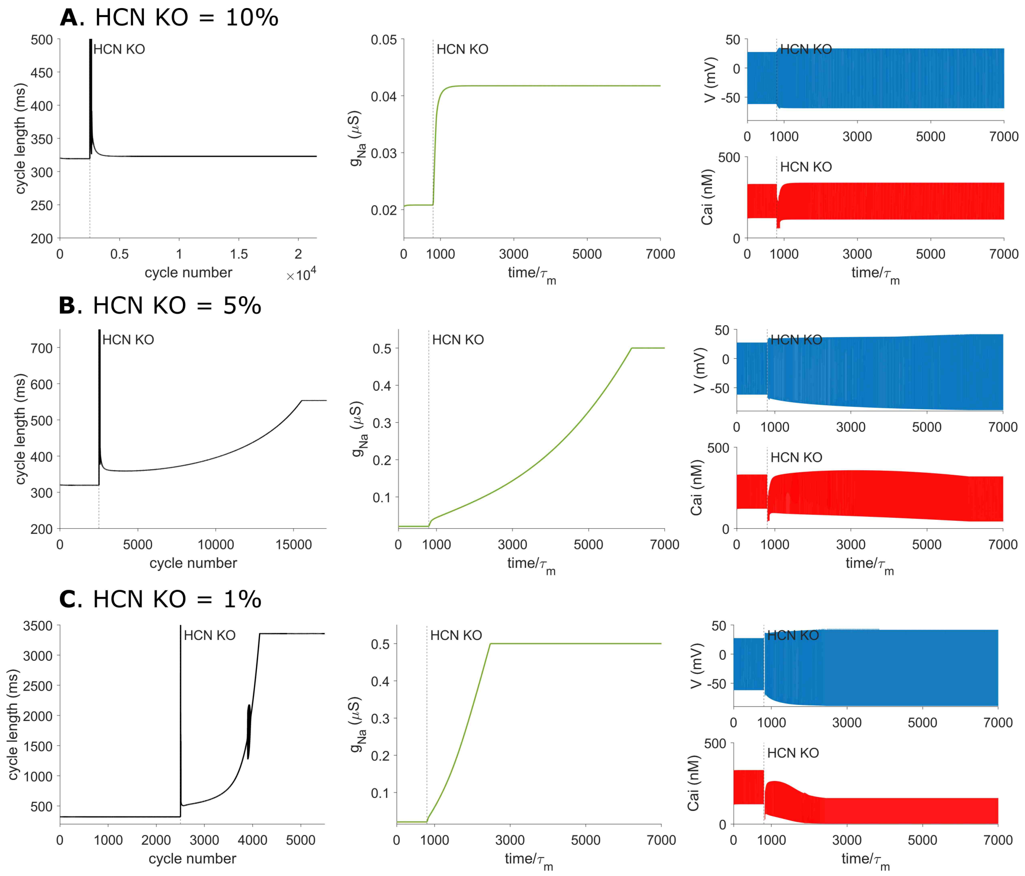


**Figure S9.** HCN KO in the single cell, for three different KO levels. Panels show from left to right: cell cycle length, $g_{Na}$, cell $V$ and $Ca_i$.

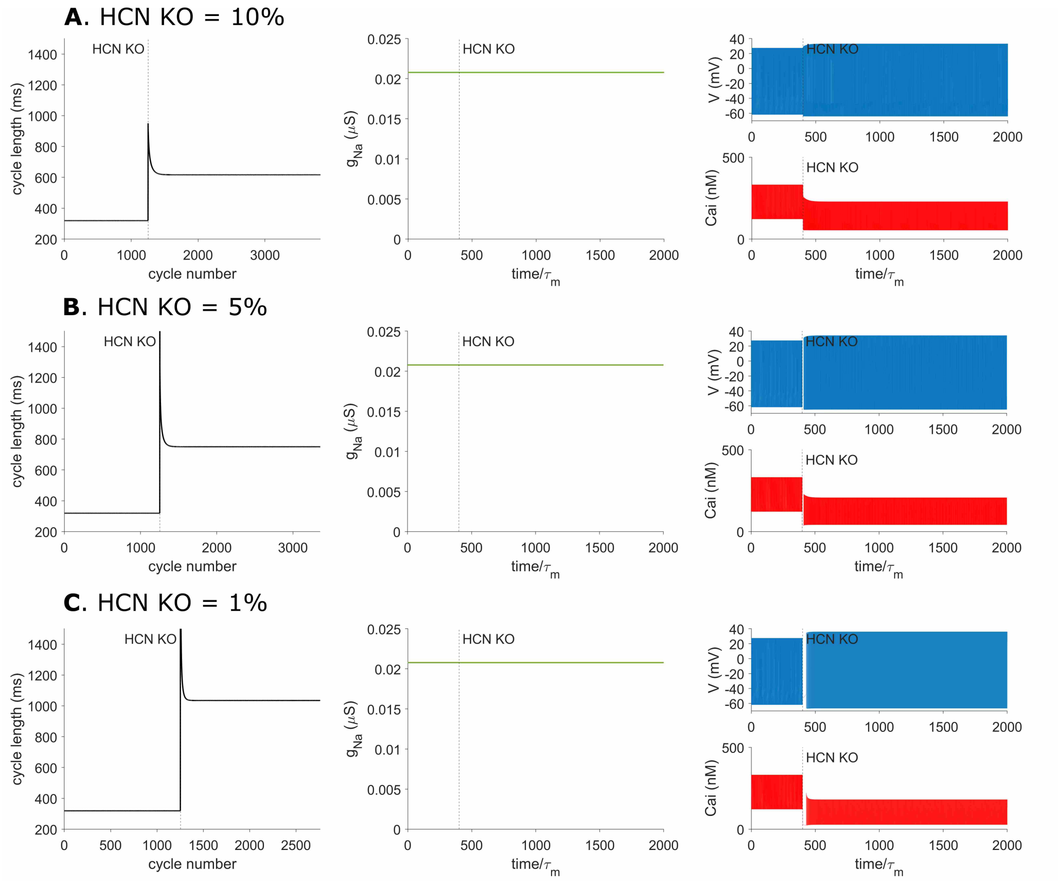


**Figure S10.** HCN KO in the single cell without feedback (fixed steady state conductances), for three different KO levels. Panels show from left to right: cell cycle length, $g_{Na}$, cell $V$ and $Ca_i$.

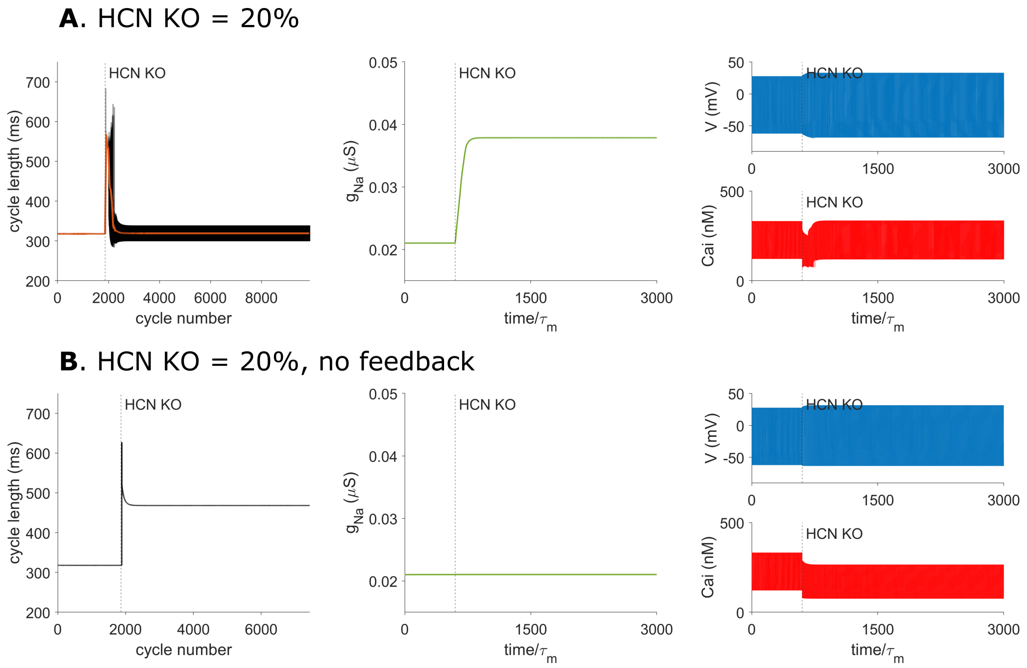


**Figure S11.** Partial HCN KO, with and without conductance feedback. Panels show, from left to right: cell cycle length (with average cycle length in red), $g_{Na}$, cell $V$ and $Ca_i$.

### Supplemental Table

**Table S1.** Calcium feedback model time constant parameters.

| Parameter | Single cell values | Tissue values | Units |
|---|---|---|---|
| $\tau_{g_{Na}}$ | 0.5 | 0.05 | μM s μ$S^{-1}$ |
| $\tau_{g_f}$ | 0.208 | 0.0208 | μM s μ$S^{-1}$ |
| $\tau_{g_{to}}$ | 3.125 | 0.3125 | μM s μ$S^{-1}$ |
| $\tau_{g_{Kr}}$ | 2.888 | 0.288 | μM s μ$S^{-1}$ |
| $\tau_{g_{Ks}}$ | 3.770 | 0.377 | μM s μ$S^{-1}$ |
| $\tau_{i_{NaK\ max}}$ | 9.92e-2 | 9.92e-3 | μM s $nA^{-1}$ |
| $\tau_{K_{NaCa}}$ | 1.55e-3 | 1.55e-4 | μM s $nA^{-1}$ |
| $\tau_{P_{CaL}}$ | 3.12e-2 | 3.12e-3 | μM s $(nA/mM)^{-1}$ |
| $\tau_{P_{CaT}}$ | 0.312 | 0.0312 | μM s $(nA/mM)^{-1}$ |
| $\tau_{P_{up\ SERCA}}$ | -5.205e-04 | -5.205e-05 | μM s $(mM/s)^{-1}$ |
| $\tau_g$ | 1 | 1 | s |

### Supplemental Movies

**Movie S1.** Time course of conductances in a 2D SAN tissue. (Left) Spatial map of sodium conductances and (Right) time course of conductances for all 400 cells in the SAN tissue.

**Movie S2.** Spontaneous activity with variable cycle length in a 2D SAN tissue. Voltage (top left) and calcium (top right) spatial maps and voltage (bottom left) and calcium (bottom right) traces for all 400 cells for the 5 different activation patterns, illustrating multiple pacemaking sites.
